# Single-molecule FRET with a minimalistic 3D-printed setup and dyes in the blue-green spectral region

**DOI:** 10.64898/2025.12.16.694555

**Authors:** Gabriel G. Moya Muñoz, Jorge R. Luna Piedra, Pazit Con, Mostofa A. Rohman, Siyu Lu, Thomas-Otavio Peulen, Thorben Cordes

## Abstract

Förster Resonance Energy Transfer (FRET) is a powerful technique for the detection and characterization of biomolecular interactions and conformational changes with sub-nanometer spatial resolution and a temporal resolution down to the timescale of fluorescence. While the technique is widely adopted in structural biology and biophysics, the evolution of single-molecule FRET has led to experimental setups with sophisticated optical layouts, multi-laser excitation schemes and time-resolved detection electronics. We here present an accessible alternative towards single-molecule FRET based on Brick-MIC, a recently introduced 3D-printed micro-spectroscopy platform. The FRET-Brick uses continuous-wave excitation at 488 nm with a minimal set of opto-mechanical components and photomultiplier detectors (PMTs). With this we were able to significantly reduce the setup complexity retaining single-molecule sensitivity with dyes matching the sensitivity of PMTs. To maximize the photon output of Alexa488, ATTO488 (donors), Alexa555, ATTO542 and Cy3B (acceptors), we introduce ferrocene-derivatives as photostabilizers that increase both dye brightness and remove dark-states. We benchmark the performance of the FRET-Brick with fluorophore-labelled oligonucleotide reference structures also in comparison to accessible volume simulations, and by detecting conformational changes in bacterial substrate binding proteins. Our work demonstrates that qualitative and quantitative smFRET measurements are possible with the minimalistic and cost-effective FRET-Brick.

## INTRODUCTION

Single-molecule Förster Resonance Energy Transfer (smFRET) has become an established technique for studying biomolecular interactions and conformational dynamics of biomolecules *in vitro* and *in vivo*^1–11^. Over time, smFRET instrumentation has evolved to include multi-laser excitation schemes, complex timing electronics, and sophisticated optical arrangements to improve its sensitivity, but also the temporal and spatial resolution^[12–15]^. Multi-parameter fluorescence detection (MFD^[16,17]^), alternating-laser excitation (ALEX^[12,18,19]^) and pulsed interleaved excitation (PIE^[13]^) have significantly advanced smFRET by enabling precise characterization of biomolecular labelling with donor- and acceptor dyes and the availability of multi-dimensional data sets and in depth information on the biomolecular system^[13,18,19]^. These allow for the correction of setup-dependent artifacts such as spectral cross-talk, differences in quantum yields between donor and acceptor dyes, and the wavelength-dependent detection efficiencies of the optical system^[6]^. Correcting for these is a requirement to convert FRET efficiency into accurate inter-dye distance (e.g., for integrative structural modelling)^[12]^ but also to identify conformational dynamics and structural ensembles. In essence, many technical developments focused on improving the sensitivity, precision and accuracy, and the information content of smFRET as a quantitative nanoscale ruler^[1,2]^.

Together with advancements in the standardization of measurement- and analysis routines, (sm)FRET is by now an established tool for mechanistic biophysical studies but also a quantitative method in structural biology^[1–11,20–28]^. The required complexity, but also the costs related to the purchase of an smFRET-instrument, raised the bar for non-specialized laboratories to enter the field and independently conduct smFRET experiments. Indeed, many applied users of smFRET have the goal to only qualitatively visualize ligand binding, domain reorientation, or conformational switching. These studies often follow an iterative process: comparing apparent FRET efficiencies under different biochemical conditions, identifying contrasts, and inferring mechanistic insights. For that purpose measurement of relative FRET efficiency changes are sufficient and extracting absolute distances (or other spectroscopic parameters such as lifetime or anisotropy) are not strictly required^[20–28]^. Thus, setups with reduced complexity would suffice for such questions since they allow to address (most) relevant aspects of the research question.

Recent efforts from our lab and others^[29–36]^ demonstrated the power of compact and affordable microscopy designs suitable for advanced optical imaging^[29,31,33–35]^. This open-source microscopy direction included applications such as smFRET^[29]^ and super-resolution microscopy^[29,34,35]^ with 3D-printed scaffolds^[29,30,33]^ and simplified optic layouts. Building on this foundation, we here present a minimalistic smFRET extension of our Brick-MIC platform^[29]^ that greatly reduces instrumental complexity and costs. The confocal FRET-BRICK is capable to detect individual diffusing molecules and uses a simple 488 nm continuous-wave (cw) excitation laser, detection via photomultiplier tubes (PMTs) and simple timing electronics. This configuration allows to conduct fluorescence correlation spectroscopy (FCS) as well as smFRET of diffusing molecules. To comply with the spectral requirements of the PMT-detectors in the setup, we introduce donor-acceptor pairs in the blue-green spectral region (Alexa488, ATTO542 and Cy3B), for which we optimized the photon output with ferrocene-based photostabilizers.

Despite all cutbacks and compromises made on the technical specifications of the FRET-Brick and the selected dye combinations, we demonstrate that the setup is suitable for the relevant mainstream biological applications of smFRET and FCS with diffusing molecules. The device allows to characterize conformational changes in a macromolecule via relative changes in FRET efficiency or differentiate distinct inter-dye distances via the use of uncorrected apparatus-dependent apparent FRET-efficiencies E*. To maximize the usage of all information contained in our single-molecule FRET data, which lacks the advanced capabilities of MFD^[16]^, PIE^[13]^ or ALEX^[18]^, we tested readily-available parameters to construct multi-dimensional FRET-histograms: the macro-time difference between donor and acceptor photons within a burst (ΔTDA) and the donor- and acceptor brightness. We found that these parameters, particularly a 2D-histogram of ΔTDA-E* enable a straightforward identification of distinct FRET-species. Using this setup and streamlined analysis approach, we demonstrate our capability of characterizing static oligonucleotide DNA standards with varying donor-acceptor distances and probing biologically relevant conformational changes in protein systems. Finally, we explored whether data from our setup can be used to obtain accurate FRET efficiency values to obtain real-space inter-dye distances for oligonucleotide standards.

## RESULTS AND DISCUSSION

### Design of the FRET-Brick

With the main goal to reduce both the costs and the complexity of confocal single-molecule detection, we chose the design of our previously published µFCS setup as a starting point for the FRET-Brick. In the developed instrument, which is described and shown in full detail in Supplementary Note 1, continuous-wave (CW) excitation is supplied by a USB-powered blue laser-diode (488-30-1235-BL, Q-LINE) and combined with detection of fluorescence from diffusing single-molecules by photomultiplier tubes (PMTs) rather than avalanche photodiodes (APDs); Figure 1. As previously described^[29]^, confocality is achieved by coupling of the emission beam into a single-mode optical fiber with a diameter of 10 µm instead of a pinhole. This simplifies the overall optical design and allows autocalibration via two piezo mirrors M5/M6 (Figure 1a). The emission is then guided to an external detection box, where it is spectrally split before reaching the detectors. This configuration is advantageous because the system functions as a lens-free microscope, apart from the objective.

**Figure 1.**
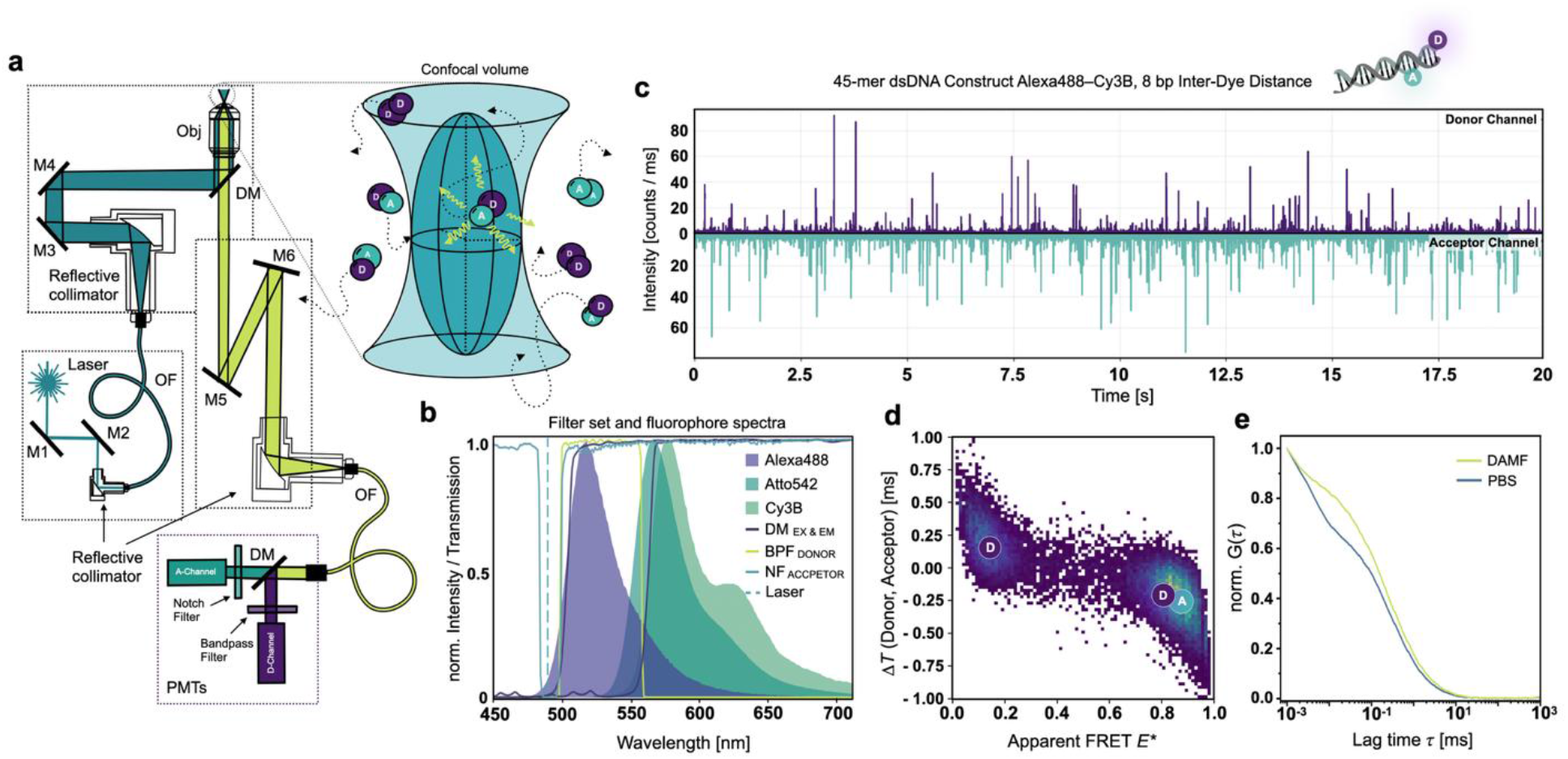
Overview of the FRET-Brick. **(a)** Optical configuration of the FRET-Brick. A 488 nm laser diode (Laser, blue) is coupled into an optical fiber (OF) and directed through the excitation layer onto the objective (Obj) to generate a confocal excitation volume. Freely diffusing, doubly-labeled molecules carrying donor (D) and acceptor (A) fluorophores emit fluorescence bursts (light green) upon entering this volume. The emitted light is collected by the same objective, coupled into an OF, and delivered to a detection module (DM) where donor and acceptor signals separated by a dichroic mirror (DM) are detected by photomultiplier tubes (PMTs). **(b)** Spectral configuration. Excitation and emission dichroic mirrors (DC) are shown in purple, the band-pass filter (BPF) for the donor channel in light green, and the notch filter (NF) for the acceptor channel in blue. The dashed blue line shows the laser excitation wavelength. Emission spectra of available fluorophores—Alexa 488 (purple), Atto 542, and Cy3B (shades of green)—are overlaid to illustrate spectral overlap with the filter set and detector sensitivity. **(c)** Representative fluorescence time trace. Twenty-second single-molecule trace of a 100 pM dsDNA sample labeled with Alexa 488 (Donor channel) and Cy3B (Acceptor channel) at 8 bp inter-dye separation, recorded simultaneously in donor and acceptor channels under 488 nm excitation at 200 µW with 100 µM DAMF ((dimethylaminomethyl)ferrocene). **(d)** (ΔT–E) histogram (apparent FRET efficiency, *E*\*, versus macro-time difference between donor and acceptor photons, Δ*T*(Donor, Acceptor)) of the dsDNA sample shown in (c), where D denotes the donor-only population and DA the FRET population. **(e)** FCS Raw data of a dsDNA sample labeled with Alexa 488, excited at 15 kW cm^−2^ in PBS buffer with and without 100 µM DAMF.

Dichroic mirrors and emission filters were selected to match the new excitation (excitation dichroic ZT491rdc, Chroma) and emission wavelengths (emission dichroic ZT543rdc, Chroma), which are in line with the higher detection efficiency of the PMTs in the blue/green spectral range (Figure 1b). To maximize the photon collection efficiency for both PMTs, a bandpass filter (FF03-525/50, Semrock) and a notch filter (NF488-15, Thorlabs) were used in the donor and acceptor channel, respectively. The latter filter blocks only reflected or scattered excitation light and and allows to collect the full acceptor emission spectrum (Figure 1b). This is particularly advantageous because the count sensitivity of PMTs decreases strongly toward longer wavelengths, dropping from about 2 × 10^5^ s^−1^ pW^−1^ at 550 nm to 1 × 10^5^ s^−1^ pW^−1^ at 600 nm and approximately 5 × 10^3^ s^−1^ pW^−1^ near 700 nm. (Figure S1). Data were collected using a USB counter module (USB-CTR04, Measurement Computing), as introduced by the Gambin lab^[30]^ and previously implemented in our FCS modality^[29,30]^, along with bespoke Python-based acquisition software capable of registering data at 1 µs time-resolution in the widely accepted photon hdf5 format^[37]^.

Based on the optical configuration shown in Figure 1, we performed smFRET experiments with excitation at 488 nm at excitation intensities of 50-200 µW using a high NA water immersion objective (60×, NA 1.2, UPlanSAPO 60XW, Olympus, Japan). Epi-fluorescence detection gave rise to count rates of both donor and acceptor dyes in the range of 10-100 kHz (Figure 1c). Thus, single-molecule transits of double-stranded 45-mer oligonucleotide structures through the confocal volume were clearly visible as burst events (Figure 1c). For the isolation of burst-data from our hdf5 data sets we used a customized version of ChiSurf^[38]^, which are available as Jupyter Notebooks with this manuscript (see Data availability statement). The software allows to obtain 1D- and 2D-histograms of apparent FRET efficiency, in which we found FRET populations to be clearly distinguished from single-donor species (Figure 1d). Similarly, the data can be used for an autocorrelation analysis which provides the typical output of an FCS experiment, i.e., diffusion- and bunching times with a temporal resolution down to ∼1 µs, only limited by the time-resolution of the used counter module (Figure 1e).

To evaluate the influence of the objective on detection performance, we compared several alternative objectives using fluorescence correlation spectroscopy (FCS). The performance was assessed via the count rate per molecule (CPM) relative to the reference 60× NA 1.2 water immersion objective (UPlanSAPO 60XW, Olympus). A 60× NA 1.35 oil immersion objective (Olympus) showed a comparable CPM (∼104% relative to the reference), as expected for a similarly high numerical aperture. In contrast, a generic low-cost OEM 100× NA 1.2 oil immersion objective and a generic OEM 60× NA 0.65 air objective exhibited substantially reduced CPM values of ∼17% and ∼1% of the reference objective, respectively (Figure S2). We further evaluated the suitability of these objectives for smFRET measurements. Representative smFRET time traces and apparent FRET efficiency histograms obtained with the three high-NA immersion objectives are shown in Figure S3. All three immersion objectives allowed detection of single-molecule burst events and determination of FRET efficiencies, with the expected reduction in photon count rates for the lower-performing objectives. Thus, smFRET measurements remain technically feasible with these objectives, albeit requiring longer acquisition times to obtain comparable statistics. In contrast, the low-NA air objective did not yield detectable burst events under the tested conditions and was therefore not suitable for smFRET measurements.

### Optimization of dye-combinations for FRET in the blue-green spectral region

Blue and green fluorescent dyes that can serve as donor and acceptor pairs in FRET experiments have a small spectral separation and a limited photostability ^[39–41]^. This poses challenges on the microscope design and the selection of matching fluorescent dyes. Based on our filter set and the dyes available in our stocks, e.g., for protein or oligonucleotide labelling, the best candidates for FRET-donors were Alexa Fluor 488 (Alexa488) and ATTO488 in combination with either Alexa555, ATTO542 or Cy3B as acceptor dyes. Each dye was characterized individually by using a 45-mer double-stranded DNA (dsDNA) sample via FCS measurements. With this we tried to identify the condition that maximized molecular brightness before the onset of photobleaching in excitation power series. We balanced molecular brightness with photo-induced bunching amplitudes observed in FCS, for instance caused by formation triplet- or dark-states. Based on our initial experiments we discarded two dyes from a detailed screening process. Alexa488 was notably brighter compared to ATTO488 (Figure S4, Table S1) despite the structural similarity of the core dye^[40,41]^ (Figure S4a). On average and independently of the conditions, Alexa488 was >2-fold brighter than ATTO488 (Figure S4b, Table S1). Consequently, all FRET-experiments were conducted with Alexa488 as donor dye. Alexa555 was also discarded from detailed analysis, since it showed much lower apparent FRET-efficiencies values compared to ATTO542- and Cy3B-labelled samples both on DNA and proteins (Figure S5) despite a reasonable burst-brightness in our experiments. For the remaining three dyes we systematically optimized combinations of donor and acceptor dye pairs, laser excitation powers, and buffer additives for improved photostability. In that regard we found it to be essential for high quality experiments in the FRET-Brick to suppress dark-state formation of the donor Alexa488 and to minimize acceptor photobleaching (Figure 2).

**Figure 2.**
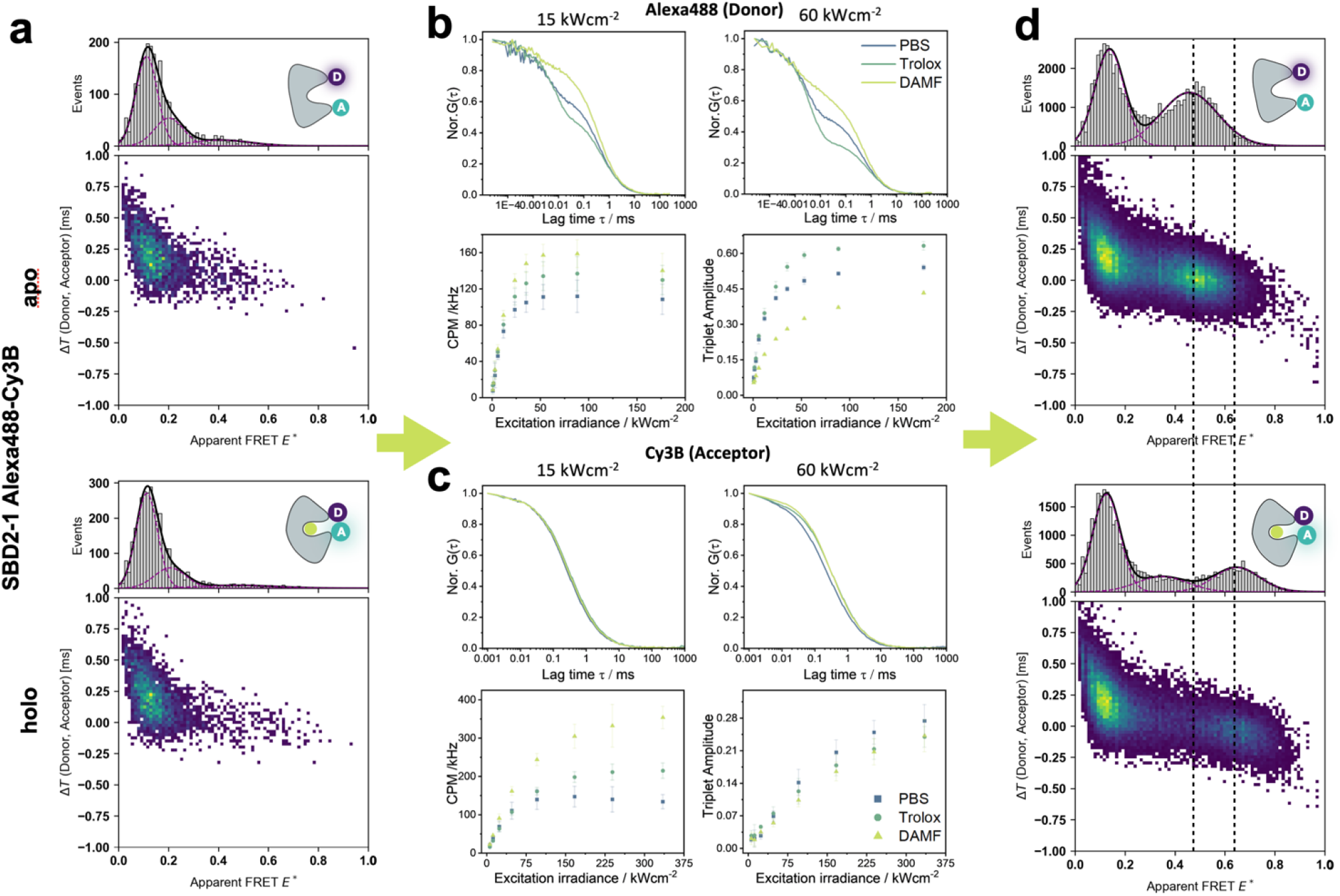
FCS-guided assay optimization and photostabilizer screening. **(a) Initial conditions of SBD2 in PBS buffer.** ΔT–E histograms of SBD2 from *Lactococcus lactis* labeled with Alexa 488 (donor) and Cy3B (acceptor), recorded at 60 kW/cm^2^ laser excitation without photostabilizing additives. The upper panel shows the open (apo) conformation, and the lower panel shows the closed (holo) conformation in the presence of 1 mM glutamine. Apart from the donor-only species, FRET-active populations are barely visible due to rapid acceptor photobleaching under these conditions. **(b**,**c) Photostabilizer screening via FCS**. Fluorescence correlation spectroscopy (FCS) of Alexa 488-labeled dsDNA (b) and Cy3B-labeled dsDNA; data for ATTO542 is presented in Figure S6 (c). Upper panels show representative normalized FCS curves at 15 kW/cm^2^ (left), and 60 kWcm^2^ (right) excitation power: blue, without additives; green, with 1 mM Trolox ((±)-6-hydroxy-2,5,7,8-tetramethylchromane-2-carboxylic acid); and yellow, with 100 µM DAMF ((dimethylaminomethyl)ferrocene). Lower panels summarize the power-dependent screening results for molecular brightness (left) and triplet-state amplitude (right), demonstrating the photostabilizing effects of Trolox and DAMF. **(d) Optimized conditions of SBD2 in PBS with photostabilizers**. ΔT–E histograms of the same SBD2-1 sample shown in (a), recorded under optimized conditions (30 kW/cm^2^ excitation, 100 µM DAMF, and 100 µM Trolox). Under these conditions, FRET-active populations are clearly visible alongside donor-only species, with the apo (open) conformation showing an apparent FRET efficiency of 0.46 and the holo (ligand-bound, closed) conformation showing E* = 0.65.

Alexa488 displayed a substantial bunching amplitude below 100 µs in PBS buffer; even at low excitation powers: >20% at 15 kW/cm^2^, ∼55% at 60 kW/cm^2^ (Figure 2b). In a FRET pair, the formation of triplet- or dark-states of the donor dye, will reduce the count-rate and increase the shot-noise of both, the donor and acceptor signal. Moreover, accurate FRET-efficiencies require dark-state corrected fluorescence quantum yields^[1,2]^. On the other hand, bleaching of the acceptor within a burst results in a bridge between the donor-acceptor and donor-only population, which complicats the determination of FRET efficiencies. To mitigate these effects, we first tested the gold-standard for photostabilization Trolox^[42]^ ((±)-6-Hydroxy-2,5,7,8-tetramethylchromane-2-carboxylic acid)^[43]^. Trolox showed a noticeable improvement of the Alexa488 count-rate (expressed in counts per molecule, CPM^**1**^). Nevertheless, it surprisingly increased the percentage of dark-state formation, particularly in the desired high excitation intensity regime >15 kW/cm^2^ (Figure 2b). In search for alternative photostabilizers, we tested ferrocene derivatives^[44]^, despite their ability for singlet-quenching of fluorescein^[45]^. (Dimethylaminomethyl)ferrocene, abbreviated DAMF, proved to be an effective additive for Alexa488. The addition of 100 µM DAMF reduced the bunching amplitude from approximately 55% to 30%. Our interpretation is that photoinduced electron transfer (PET^[46]^) occurs between the triplet-state of Alexa488 and DAMF resulting in the formation of a radical anion from which the singlet ground-state can be recovered by molecular oxygen. This ROXS-process^[47]^ increased the molecular brightness of Alexa 488 by ∼1.5-fold to maximum values of 160 kHz at excitation powers >60 kW/cm^2^.

Strikingly, the effects of DAMF on the performance of Cy3B (Figure 2c) and ATTO542 (Figure S6) were similar and again DAMF outperformed Trolox. DAMF showed 1.5-3-fold increases in CPM with a similar decrease in bunching amplitudes (Figure 2c). We were surprised about the competitive performance of DAMF and will present further results of similar stabilizers in upcoming work. Based on the optimized conditions for excitation intensity and photostabilizer conditions (100 µM DAMF) we performed FRET experiments with both combinations of donor and acceptor dyes on substrate-binding domain 2 (SBD2), the soluble extra-cellular component of an amino-acid import system of *Lactococcus Lactis*^[48]^. Without any detailed analysis, the differences of the histogram quality between PBS buffer with (Figure 2d) and without the photostabilizer DAMF (Figure 2a) are strikingly clear and support the relevance of our screening efforts for optimized photostability.

### smFRET Measurements with the FRET-Brick

Using the optimized conditions, we performed FRET measurements on a series of double-stranded DNA (dsDNA) and SBD2 protein samples to evaluate the performance and dynamic range of the FRET-Brick. Initial reference measurements were carried out with 45-mer dsDNA constructs^[49,50]^ with the dye pair Alexa488-Cy3B. The experiments included donor-only, acceptor-only, and doubly-labelled dsDNA with distinct inter-dye distance of 8 and 18 bp, as well as a mixed sample containing both 8 and 18 bp oligonucleotides (Figure 3 and Figure S7). The data were analyzed by initial burst identification and burst-wise extraction of different parameters. These were burst-wise proximity ratios to determine apparent FRET efficiencies E*, the macro-time difference ΔTDA between donor and acceptor photons within a burst and donor- and acceptor count-rates.

**Figure 3.**
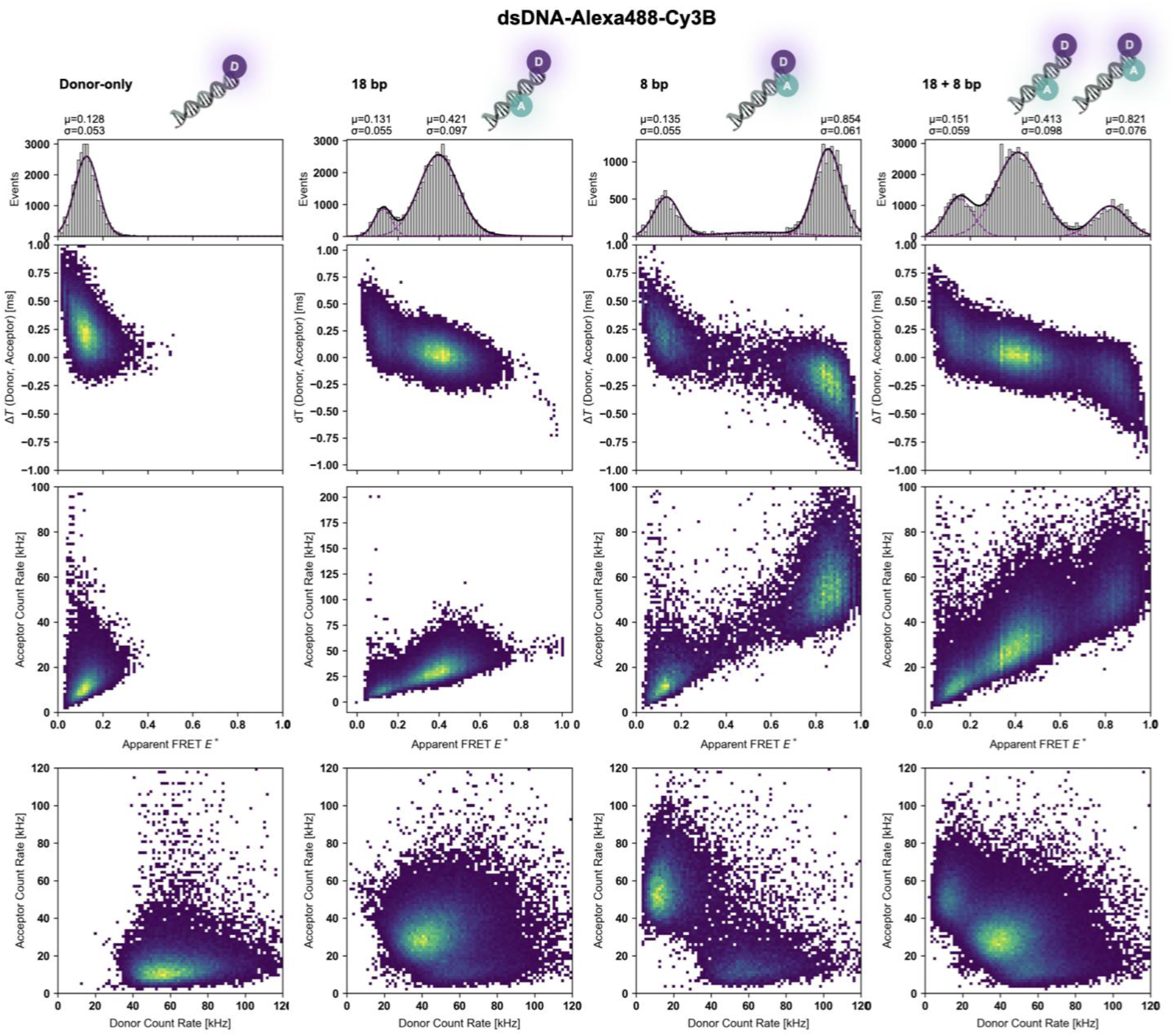
Multiparameter 2D histograms of Alexa 488–Cy3B labeled dsDNA samples. Two-dimensional histograms of a 45-mer dsDNA labeled with Alexa 488 (donor) and Cy3B (acceptor) at 60 kW/cm^2^ excitation in the presence of 100 µM DAMF: donor-only, donor-acceptor dsDNA with 18 bp, 8 bp interdye distances and a mix of the two. Rows correspond to different parameter spaces: (top) apparent FRET efficiency versus DT_DA_, (middle) apparent FRET efficiency versus acceptor brightness, and (bottom) acceptor versus donor brightness.

ΔTDA represents the macro-time difference between donor and acceptor photons within a burst, calculated from photon counts binned in 1 µs intervals. Because each bin can contain multiple photons from both donor and acceptor channels, ΔTDA does not reflect exact photon arrival times. Instead, it provides a coarse-grained, statistical measure of the relative distribution of donor and acceptor photons within the burst. Values near zero indicate that donor and acceptor photons frequently appear within the same bins, positive values are related to bins dominated by donor photons, and negative values are bins dominated by acceptor photons. Donor and acceptor brightness reflect the total photon counts per molecule in each channel and vary due to FRET efficiency in the different samples.

From these parameters, we generated a series of two-dimensional histograms to visualize population distributions with the goal of separating FRET-active molecules from donor-only species. The representations included apparent FRET efficiency E* versus ΔTDA (ΔT-E histogram), E* versus acceptor brightness (A-E histogram), and donor versus acceptor count-rate (D-A histogram); see different rows of Figure 3. In the donor-only sample, the apparent FRET efficiency was 0.13, consistent with the expected donor leakage (lk). Overall, the donor-only population was clearly distinguishable in all three plots. The acceptor-only sample did not yield a measurable signal corresponding to the acceptor dye (Figure S7). Facilitated by the 2D-plots we observed that dsDNA with both donor- and acceptor dye are readily distinguishable from donor-only species. The 18 bp and 8 bp samples had E* values of 0.42 and 0.85, respectively, giving rise to a large dynamic distance range considering the Δbp of 10. Consequently, even subtle changes in inter-dye separation are clearly resolved. In the mixed 8 bp / 18 bp sample, all three populations—donor-only, 18 bp, and 8 bp—were resolved across all plots, further demonstrating the power of the multiparameter analysis for distinguishing coexisting species.

To assess the system’s performance with a different dye pair, we studied the same dsDNA constructs with ATTO542 as acceptor dye (Figure 4) with inter-dye separations of 23 bp, 18 bp, and 8 bp. All samples had varying E*-values according to their inter-dye separation and contained a clearly visible donor-only species, which was straightforward to isolate in the 8 bp case. The apparent FRET efficiencies E* for this dye pair were 0.32 for the 23 bp construct, 0.40 for the 18 bp construct, and 0.65 for the 8 bp construct giving rise to a smaller dynamic range, either due to a different R0-value or enhanced photobleaching of ATTO542 compared to Cy3B. It was notable that some data sets (18 bp, 23 bp) contained subtle high-FRET sub-populations, which have also been reported with this dsDNA construct with other fluorophores before^[50]^, but might also be associated with small contamination of the sample with the 8 bp construct. In our view this highlights the sensitivity of our analysis for detecting even minor coexisting species. Despite these minor contaminants, all main FRET-active populations are clearly resolved, demonstrating that the system can reliably distinguish multiple constructs and effectively separate donor-only and FRET-active species across dye pairs with varying dynamic ranges.

**Figure 4.**
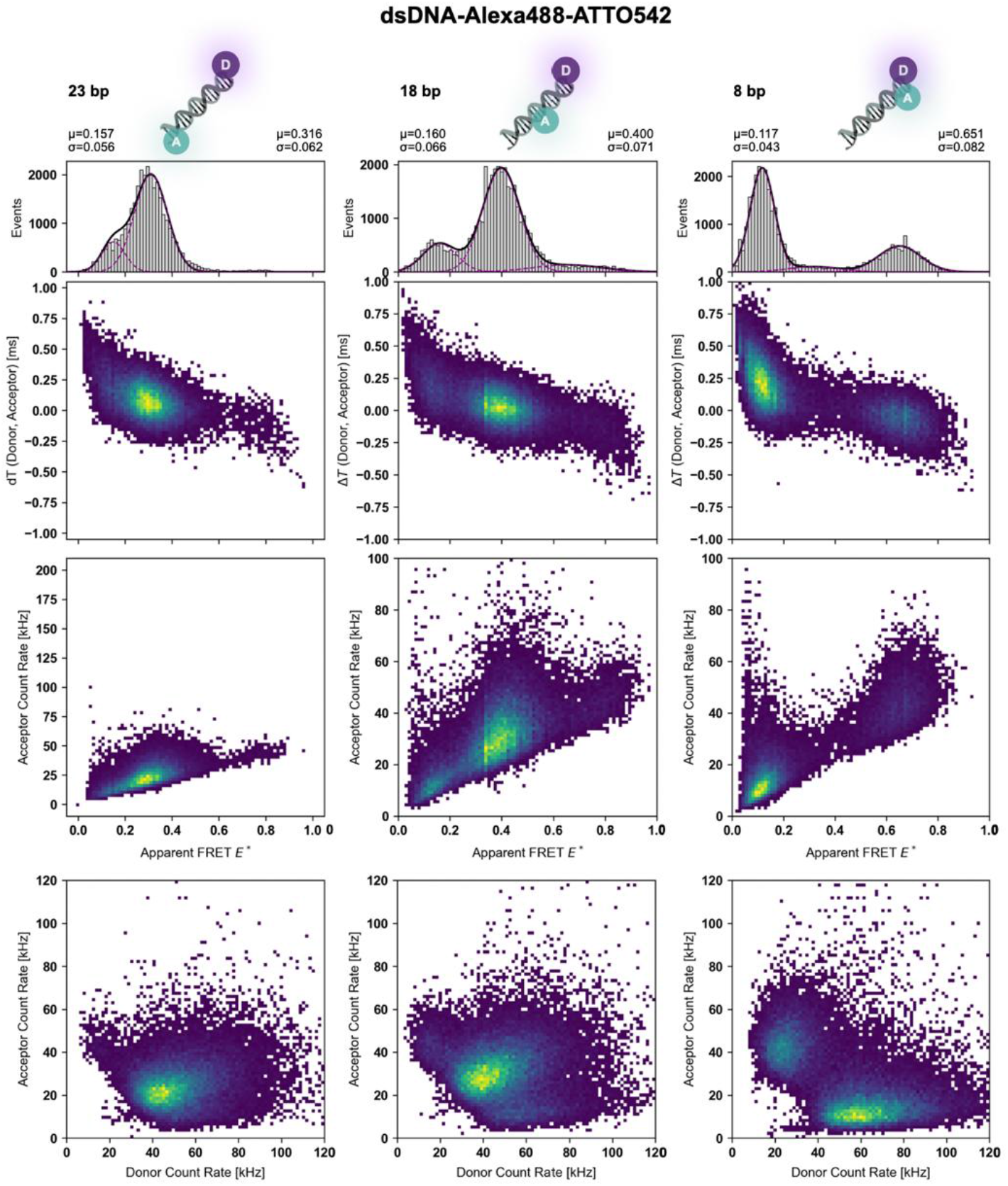
Multiparameter 2D histograms of Alexa488–ATTO542 labeled dsDNA samples. Two-dimensional histograms of a 45-mer dsDNA labeled with Alexa488 (donor) and ATTO542 (acceptor) at 60 kW/cm^2^ excitation in the presence of 100 µM DAMF. In each dataset for different donor-acceptor inter-dye distances, donor-only species are clearly visible. Rows show different parameter spaces: (top) apparent FRET efficiency versus DT_DA_, (middle) apparent FRET efficiency versus acceptor brightness, and (bottom) acceptor versus donor brightness.

Finally, to demonstrate the technical abilities of the FRET-Brick beyond static DNA samples, FRET measurements were performed on SBD2^[48]^. The protein adopts the well-established periplasmic binding protein fold^[51,52]^ in which proteins undergoes large conformational change upon ligand binding, transitioning from an open (apo) to a closed (holo) state^[26,28]^. For the smFRET experiments, we used a published mutant in which the labeling sites (369C/ 451C)^[27,53]^ were positioned such that the FRET efficiency increases when the protein adopts the holo conformation upon ligand binding (Figure 5). Initial experiments with Alexa 488 and Cy3B on SBD2 showed elevated acceptor photobleaching even with DAMF added, which was not observed to the same extent in DNA samples. This is evident in the histograms as a bridge population connecting the donor-only population and the donor-acceptor species (Figure S8). To mitigate this effect, the excitation power was reduced to 30 kW/cm^2^ and 100 µM Trolox was added additionally to stabilize the acceptor Cy3B (Figure 5 and Figure S9). Despite a notable improvement, a clear bridge population persisted (compare data in Figure S8 and S9).

**Figure 5.**
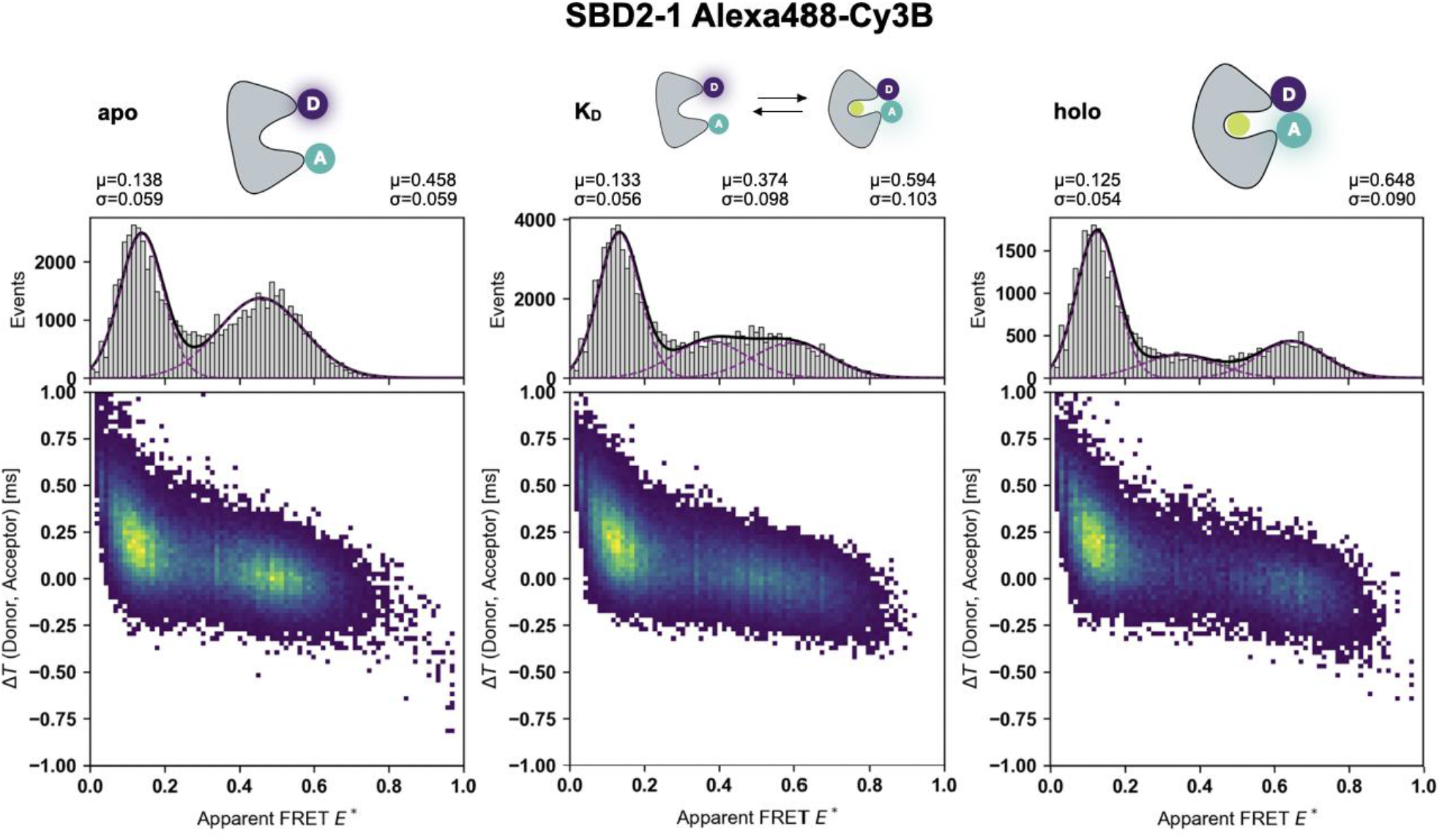
Multiparameter 2D histograms of SBD2 in different ligand-bound states with photostabilizer DAMF and Trolox. ΔT-E histograms of SBD2 with Alexa 488 (donor) and Cy3B (acceptor) at 60 kW/cm^2^ excitation with 100 µM DAMF and 100 µM Trolox. The left panel shows the apo (open), the middle and right panel shows the K_d_ (2 µM glutamine) conditions and holo state (1 mM glutamine), respectively.

It is worth noting that ΔT–E, A–E, and A–D histograms revealed FRET species that were well-separated from the donor-only peak at E* ∼0.14 (Figure 5, apo) and allowed a clear identification of photophysical processes. Based on the structure of SBD2 and previous publications, we clearly attribute the population at E* = 0.46 to the open conformation (E* = 0.46), which is dominant in the absence of ligand. Ligand addition induces a shift toward higher FRET efficiency (E* = 0.65) and represents the closed holo state of the protein adopted in the presence of glutamine. Both populations are observed at ligand conditions of 2 µM glutamine reflecting equal occupancy of the two conformational states. While this supports the idea of a functional protein, we observed that photobleaching was stronger in the holo-state reducing the overall holo-populations relative to the donor-only (Figure 5). Under such conditions determination of affinity values such as Kd via a titration will be systematically biased towards higher values, since more ligand is (apparently) needed to increase the fraction of the ligand-bound holo state.

### Benchmarking of accurate FRET and calculation of inter-dye distances

As shown in the preceding sections, the FRET-Brick and the established analysis procedure via ΔT–E* and brightness-based histograms faithfully reproduce the Förster relation qualitatively for dsDNA and ligand-induced conformational changes in SBD2. We finally wondered whether our approach and the information accessible are even sufficient to obtain accurate FRET efficiency values and interprobe distances. To benchmark our instrument, we use Photon distribution analysis^[54]^, a method that explicitly considers the photon shot-noise, to maximize the attainable precision. We developed and model that accurately accounts for the excitation and emission crosstalks (Supplementary Note 2, eqs. 3-11), implemented the model in the open-source software ChiSurf^[38]^, and applied the model to dsDNA data to benchmark our instrument and test the ability to resolve species. The experimental histogram of the apparent FRET efficiencies E* of a mix of dsDNA with 8 bp/18 bp inter-dye distances of Alexa488/Cy3B was described by a Gaussian mixture model normally distributed in the inter dye distances (Figure 6). In the analysis the mean inter-dye distances, the population fractions, the distribution width, the fraction of FRET inactive species together with the spectral crosstalk from to donor to the acceptor channel were considered as variable parameters.

**Figure 6.**
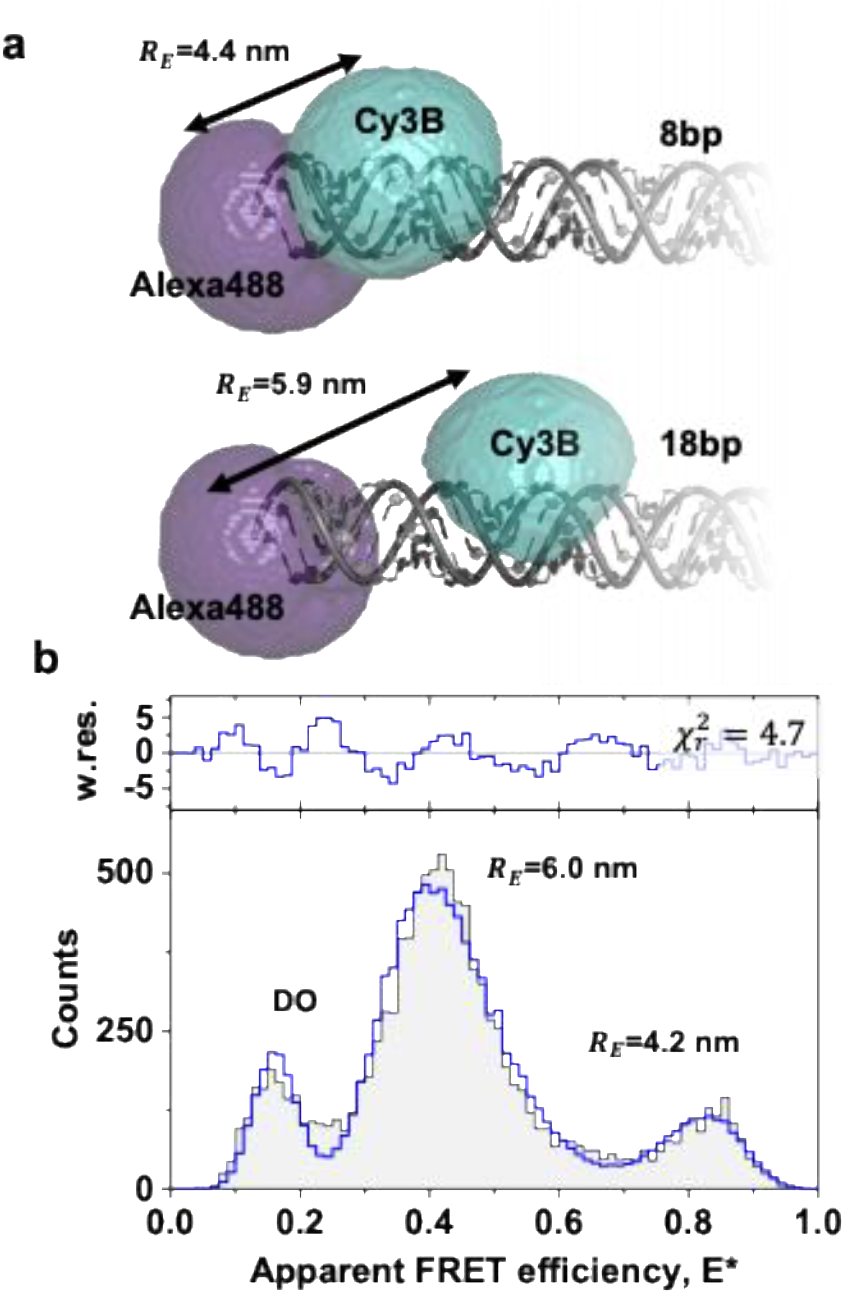
Simulated inter-dye distance and photon distribution analysis. **a)** Accessible volume simulations for Alexa488 and Cy3B attached to dsDNA separated by 8 and 18 base pairs (bp). The distances, *R*_*E*_, correspond to FRET averaged distances. **b)** Photon distribution analysis of a 8 bp and 18 bp Alexa488/Cy3B mixture. In the analysis bursts were binned in 3 ms time-windows (TWs), the photons recorded for the two detection channels were binned in a 2D histogram, which is marginalized to the apparent FRET efficiency, *E*_*app*_ (black line, gray area). The data was described by a mixture model with two normal distributions (SI Note 2, eq. 3) and a fraction of FRET inactive species (DO). The reported distances correspond the mean recovered distances. The spectral sensitives and excitation cross-sections were determined by spectra, the emission cross talk of the donor in the acceptor channel was a free fitting parameter with a fit value of 0.156). To recover absolute distances, the manufacturer provided fluorescence quantum yield of Alexa488 (Φ_*F,D*_ = 0.95) and a reference value for Cy3B (Φ_*F,A*_ = 0.85)^[55]^ were corrected by *b*_*a*_, the power induced dark state population (Φ_*sm*_ = (1 ™ *b*_*a*_)Φ_*F*_). Absolute distances were determined using a spectrum derived^[56]^ Förster radius (*R*_0_ = 6.5 *nm*) and the dark-state corrected quantum yields (Φ_*D,sm*_ ≈ 0.8, Φ_*A,sm*_ ≈ 0.65).

The spectral experimental correction parameters (see Figure 1b), i.e., the spectral excitation and emission cross talks and the spectral sensitivities of the detectors for the dyes, were computed based on public (Cy3B) data^[55,56]^ or data provided by the manufacturer (ATTO542, Alexa488). For Alexa488 the extinction coefficient at the used excitation wavelength of 488 nm was ∼75.000 L·mol^−1^·cm^−1^. At 488nm the direct excitation for ATTO542 and Cy3B for the corresponding extinction coefficients of ∼13.000 L·mol^−1^·cm^−1^ and ∼28.600 L·mol^−1^·cm^−1^, suggesting considerable excitation cross-talks, *α*_*Exc*_ = *σ*_*A*|*G*_ /*σ*_*D*|*G*_, of 0.22 (Alexa488/Atto542) and 0.48 (Alexa488/Cy3B). Based on the Alexa488 emission spectrum and the optical elements used in our setup we anticipate a donor to acceptor crosstalk (fraction of donor emission detected in acceptor channel, *α*) of 0.096. Meanwhile, based on the spectra, we anticipate donor to acceptor crosstalk of 0.021 and 0.113 for Cy3B and ATTO542, respectively. This highlights the need to account for spectral crosstalk and sensitivities for accurate FRET analysis. Based on the spectral forms, the detection efficiency ratio of the FRET pair Alexa488/Cy3B and Alexa488/Atto543 were 2.21 and 2.29, respectively. Summing up, Alexa488/Cy3B has a lower emission crosstalk, yet a higher excitation crosstalk compared to Alexa488/ATTO542.

In the past, dsDNA served as standard in international round-robin tests that showed an excellent agreement between FRET distanced modeled by accessible volumes (AVs)^[8]^ and experiments^[2]^. Thus, we simulated the FRET-averaged distances, *R*_*E*_, between the labeling sites by using AV simulations (Figure 6a). The FRET distances based on AVs simulations with the experimentally recovered distances showed a good agreement (Figure 6b). Moreover, a comparison of the optimized emission cross talk (0.15) with the cross talk expected by the spectra (0.11) is consistent supporting the idea that the FRET-Brick can be used not only for qualitative but also quantitative FRET studies.

## CONCLUSION

While innovation in single-molecule methods is often associated with increasing technical sophistication, i.e., multi-colour laser excitation^[14,57,58]^ state-of-the-art timing and detection electronics^[13,59,60]^, complex optics^[61]^ or automated sample handling^[62–64]^ this progress also comes with higher cost and complexity, unintentionally widening the accessibility gap. Innovation, however, can also go into the opposite direction: reducing a system to its essential components to make it accessible to a broader range of users. This is particularly relevant as single-molecule fluorescence approaches have become increasingly popular in biology and biochemistry.

Here, we demonstrate that single-molecule FRET can be performed with a minimal setup with auto-alignment capabilities using a laser pointer for excitation, PMTs for detection with a minimal set of optics and simple detection electronics. Rather than relying on multi-colour excitation methods such as ALEX or PIE, we extracted meaningful information beyond the ratiometric FRET-efficiency using basic burst-parameters such as donor-acceptor count rates and macrotime differences between donor- and acceptor photons. In the respective 2D-histograms we were able to identify FRET-active molecules and to distinguish them from donor-only species. With this we were able to resolve distinct FRET efficiencies in static DNA constructs. With this we obtained a good sense of the dynamic range of our instrument with different FRET pairs and were able to assess the conformational changes in a protein system. PDA and accurate FRET analysis even showed that the system was not even restricted to pure qualitative experiments of the FRET ruler (e.g., determination of relative distance changes), but that the data contained sufficient information to convert setup-dependent E* values into inter-dye distances for oligonucleotide structures. Besides all technical aspects of our work, we also introduced a new compound for photostabilization of dye molecules in the blue-green spectral region. Using the ferrocene-based compound DAMF we were able to significantly suppress dark-state formation in both donor and acceptor dyes with a simultaneous increase fluorophore brightness. The use of this photostabilizer led to markedly better FRET histogram quality for both DNA and protein samples and the success also motivated us to further study these molecules for other applications.

While we see room for improvement in both instrumentation and experimental conditions, this work serves as a proof of concept that reliable single-molecule FRET measurements can be achieved with a simple and robust optical setup. The photomultiplier tubes used here were not optimized for the FRET-assay but were simply available in the laboratory; detectors and optical layouts better suited for this spectral region could further enhance performance. Similarly, optimization of the detection volume e.g. confocal volume, such as operating beyond the diffraction-limited regime and adjusting pinhole sizes ^[1]^, has also the potential to improve data quality. Finally, assay-specific refinements, particularly for protein samples but also in the data analysis via open-source code (ChiSurf), can further extend the applicability of our method.

Together, our instrument and the obtained results highlight that meaningful, quantitative single-molecule insights can be obtained with an accessible, low-cost apparatus. Simplifying the experimental framework while leveraging digital data processing offers a path toward making smFRET more broadly available across disciplines, bridging the gap between technical innovation and practical usability.

## Acknowledgements

This work was generously supported by start-up funding of TU Dortmund University, the Bundesministerium für Bildung und Forschung (KMU grant „quantumFRET” to T.C.) and Deutsche Forschungsgemeinschaft (DFG-CO879-6-1, project Linker to T.C.). P.C. acknowledges a postdoctoral fellowship from the Alexander von Humboldt foundation.

## Author contributions statement

G.G.M. and T.C. designed and conceived the study. G.G.M., J.L.P. and T.C. built the setup. M.R. and S.L. provided data on photo-stabilizers. G.G.M., J.L.P. and P.C. conducted experiments. T.P. wrote data analysis software. G.G.M., P.C., M.R. and T.P. analyzed data. T.C. acquired funding and supervised the study. G.G.M. prepared figures and wrote the initial draft of the manuscript together with T.C. The manuscript was reviewed and approved by all authors.

## Material and Methods

Material and Methods are provided in the Supplementary Information in Supplementary Note 1.

## Data availability

The acquired experimental data and analysis software are accessible from Zenodo: 10.5281/zenodo.17921943.

## Additional information

G.G.M. and T.C. declare commercial interest in Brick-MIC as scientific co-founders of FluoroBrick Solutions GmbH a company that distributes micro-spectroscopy setups.

## Supplementary Information

**Figure S1:**
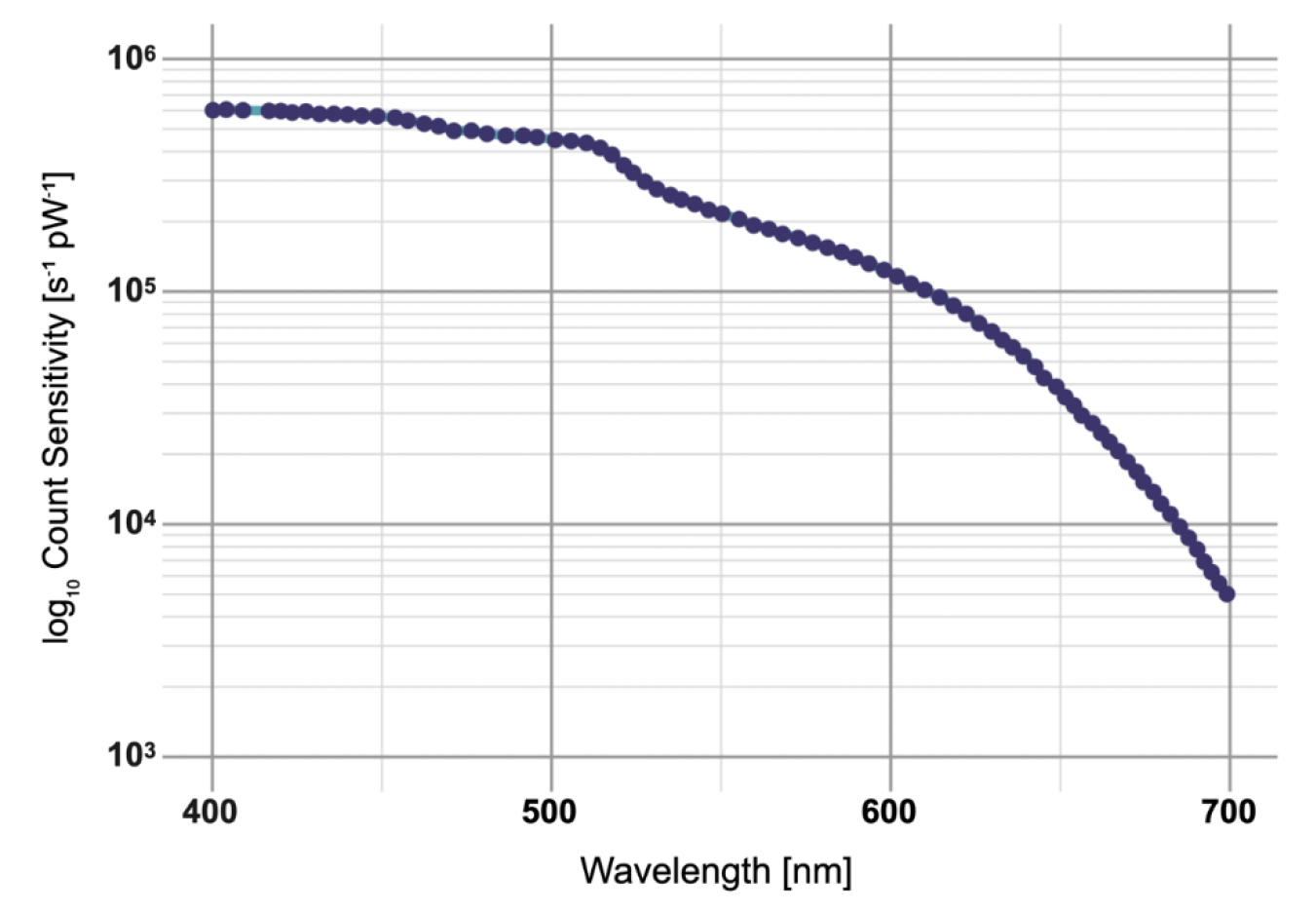
Wavelength-dependent photon count sensitivity of the Hamamatsu H10682-210 photomultiplier tube (PMT). Data extracted from the manufacturer’s specifications.

**Figure S2:**
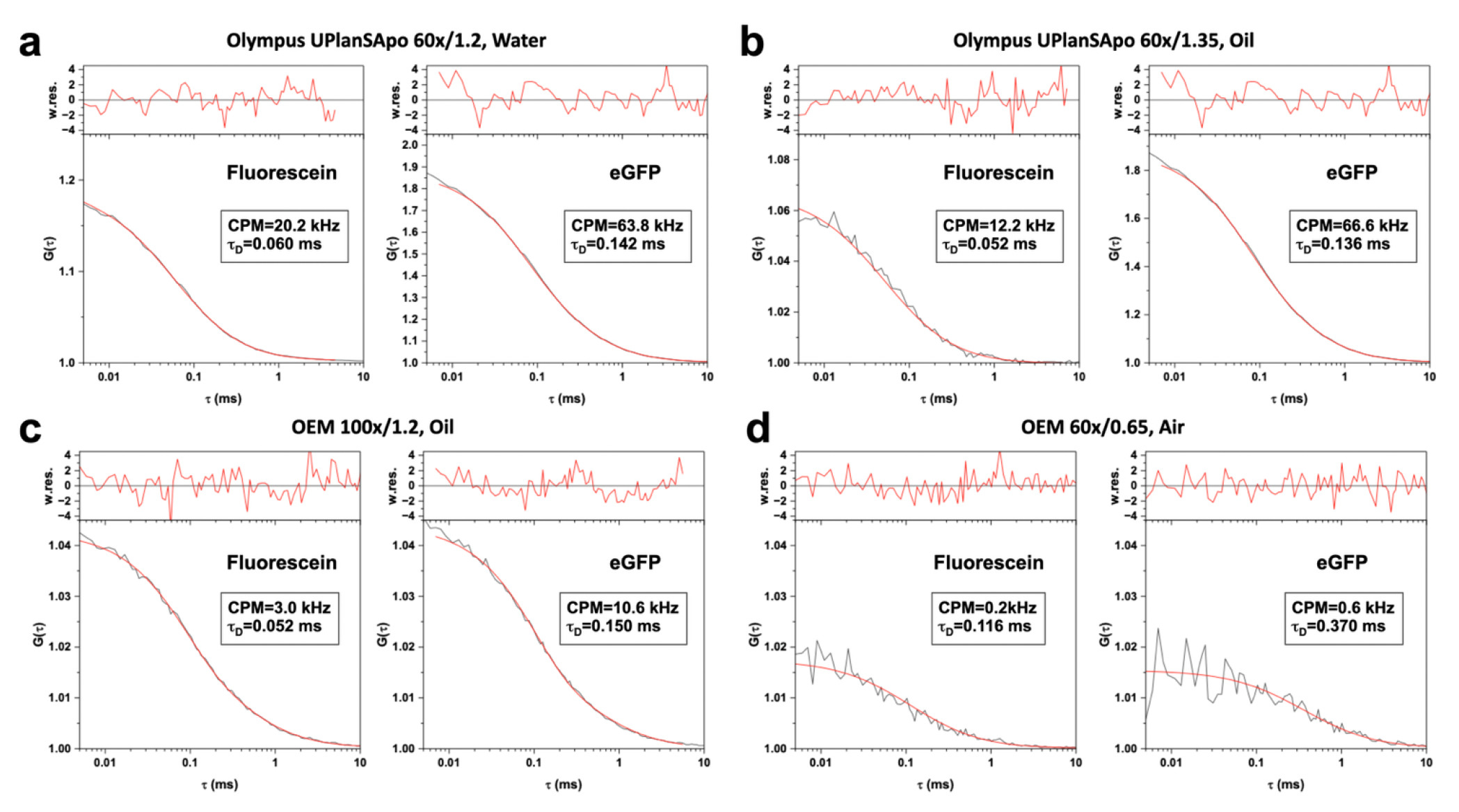
Objective-dependent fluorescence correlation spectroscopy (FCS) measurements. Autocorrelation curves recorded for fluorescein and eGFP using four different microscope objectives at an excitation power of 100 µW (measured after the objective). Panels correspond to (a) 60× NA 1.2 water immersion objective (Olympus), (b) 60× NA 1.3 oil immersion objective (Olympus), (c) generic OEM 100× NA 1.2 oil immersion objective, and (d) generic OEM 60× NA 0.65 air objective. Experimental data are shown as black curves, while the corresponding fits are shown in red.

**Figure S3:**
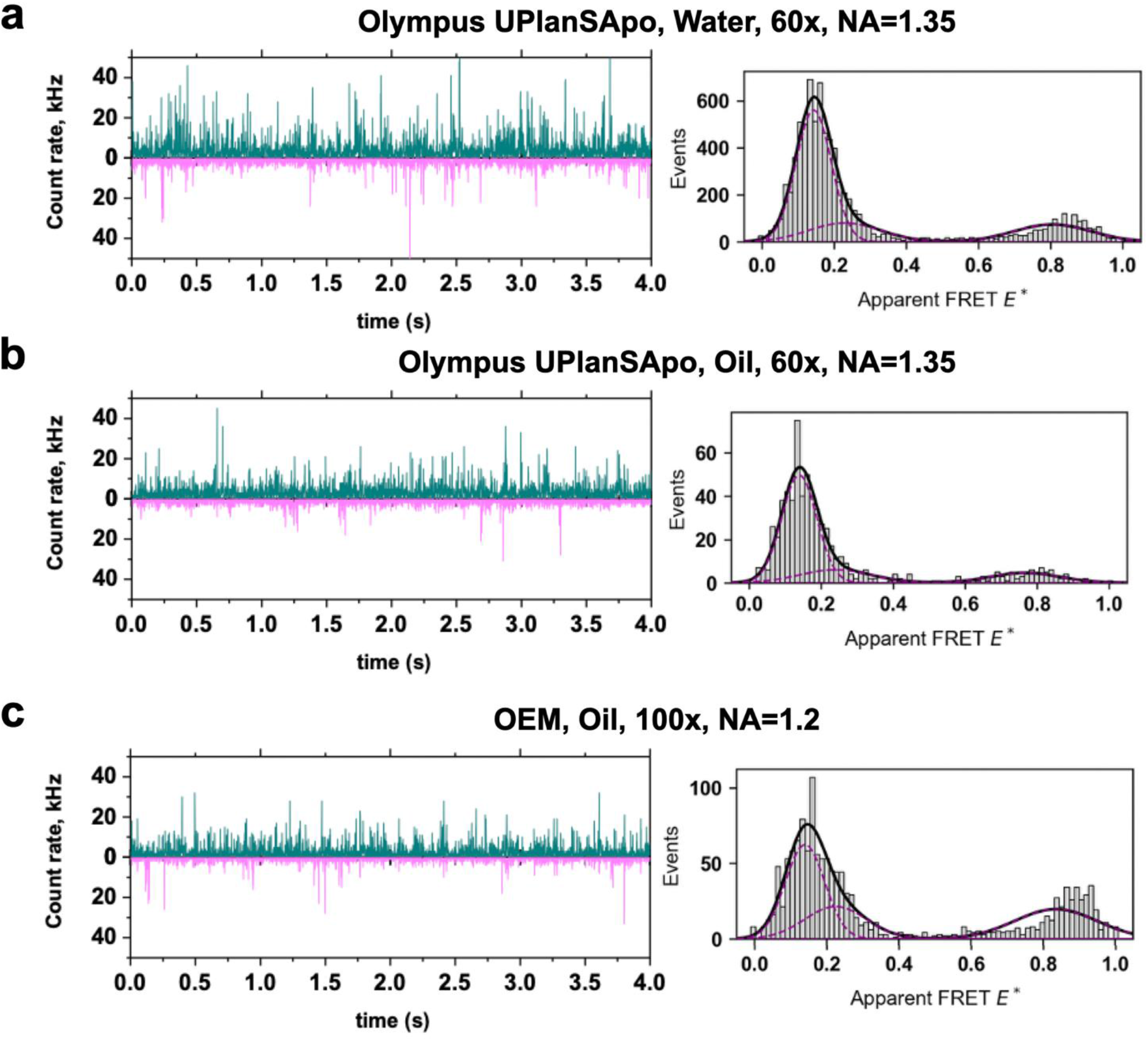
smFRET measurements obtained with different microscope objectives. Representative single-molecule time traces (left, 4 s duration) and corresponding apparent FRET efficiency histograms (right) recorded for a 45-mer dsDNA labeled with Alexa 488 (donor) and Cy3B (acceptor) with an 8 bp interdye separation. Measurements were performed using (a) a 60× NA 1.2 water immersion objective (Olympus), (b) a 60× NA 1.3 oil immersion objective (Olympus), and (c) a generic OEM 100× NA 1.2 oil immersion objective. All measurements were conducted at an excitation power of 100 µW (measured after the objective) in the presence of 100 µM DAMF. Apparent FRET efficiency histograms were obtained from 5 min measurements.

**Figure S4:**
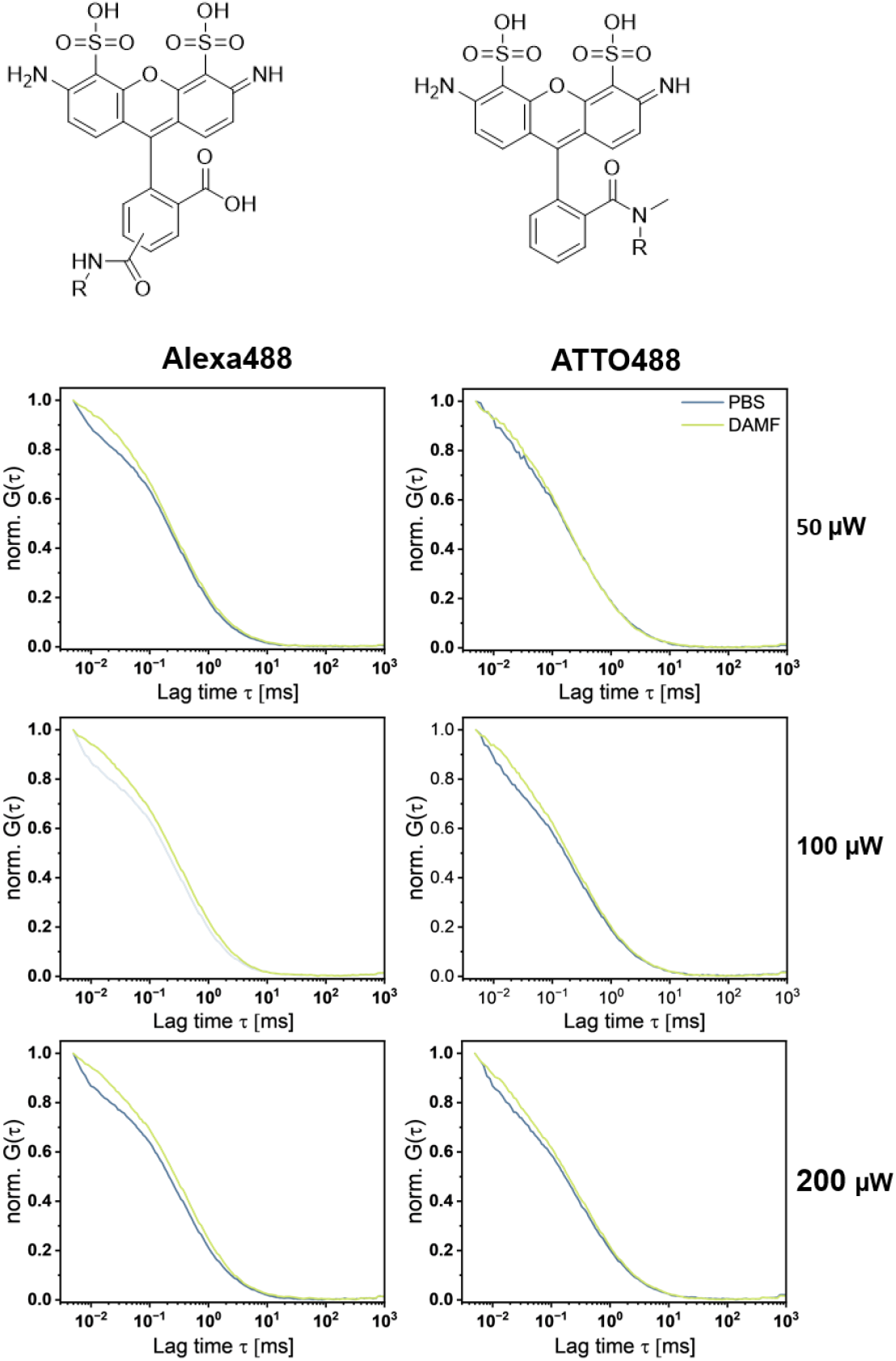
**a)** Chemical structure of fluorophores Alexa 488 (left) and ATTO488 (right). **b)** FCS power dependence of dsDNA labelled with Alexa488 (left) and ATTO488 (right), excited at 488 nm with 50 µW (top), 100 µW (middle), and 200 µW (bottom). Measurements were performed in buffer only (blue) and in the presence of 100 µM DAMF (yellow) on the FRET-Brick.

**Table S1:** **a)** Fit parameters extracted from FCS data recorded on the FRET-Brick shown in Figure S1. t_b_ = Bunching time; b_a_ = Bunching amplitude; CPM = Counts per molecule.

| Power / $\mu\text{W}$ | $\tau_b$ [$\mu\text{s}$] | $b_a$ | CPM [kHz] | $\tau_b$ [$\mu\text{s}$] | $b_a$ | CPM [kHz] |
| --- | --- | --- | --- | --- | --- | --- |
| **Alexa488** | **PBS** |  |  | **DAMF** |  |  |
| **50** | 5.6 | 0.33 | 21.6 | 19.6 | 0.19 | 19.3 |
| **100** | 5.6 | 0.37 | 21.6 | 19.6 | 0.18 | 22.1 |
| **200** | 5.6 | 0.40 | 18.5 | 19.6 | 0.19 | 22.0 |
| **ATTO488** |  |  |  |  |  |  |
| **50** | 13.0 | 0.34 | 6.9 | 36.0 | 0.27 | 7.0 |
| **100** | 13.0 | 0.36 | 9.0 | 36.0 | 0.28 | 10.8 |
| **200** | 13.0 | 0.37 | 9.9 | 36.0 | 0.30 | 12.1 |

**Figure S5:**
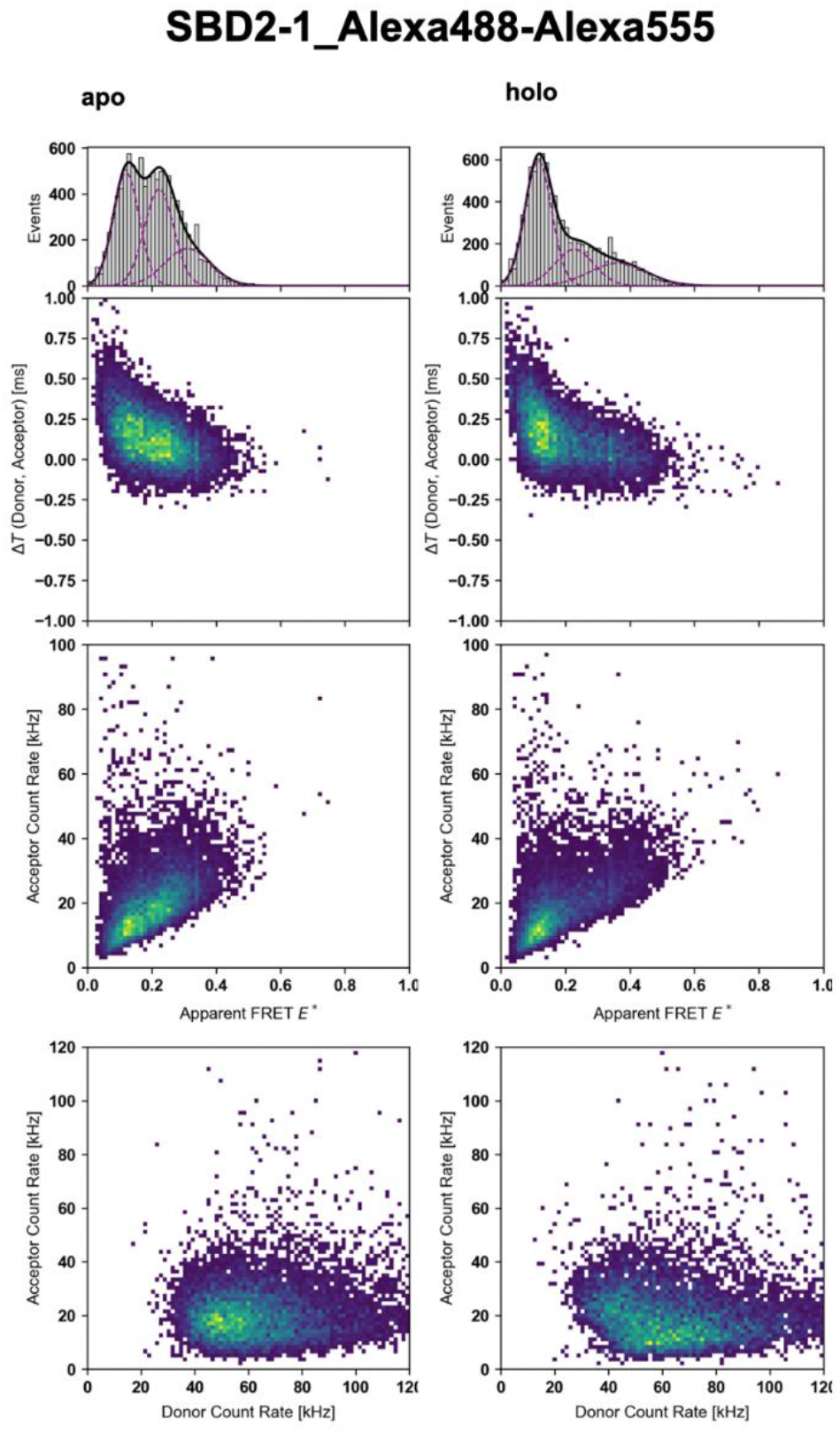
ΔT-E, A-E*, A-D histograms of SBD2 labelled with Alexa 488 (donor) and Alexa555 (acceptor) recorded at 60 kW/cm^2^ excitation power with 100 µM DAMF. The left panel shows the apo (open) conformation, the right panel shows the holo (closed) conformation in the presence of 1 mM glutamine.

**Figure S6:**
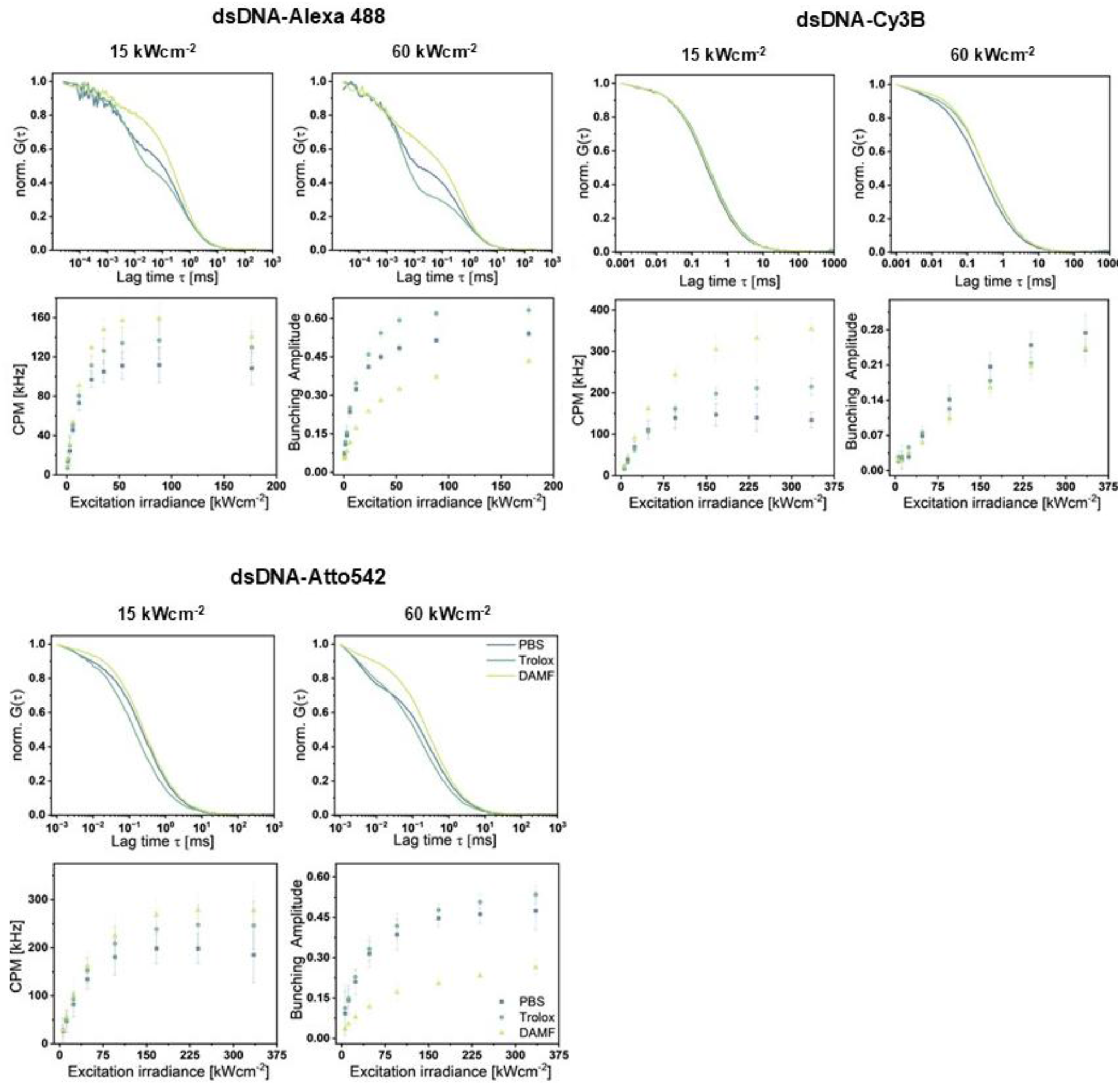
Photostabilizer screening via FCS of Alexa488, Cy3B and Atto542-labeled dsDNA. Upper panels show representative normalized FCS curves at 15 kW/cm^2^ (left) and 60 kW/cm^2^ (right) excitation power. Buffer conditions are colour-coded: blue, without additives; green, with 1 mM Trolox ((±)-6-hydroxy-2,5,7,8-tetramethylchromane-2-carboxylic acid); and yellow, with 100 µM DAMF ((dimethylaminomethyl)ferrocene). The lower panels summarize the power-dependent screening results for molecular brightness (left) and bunching amplitude (right) to validate the positive photostabilizing effects of Trolox and DAMF.

**Figure S7:**
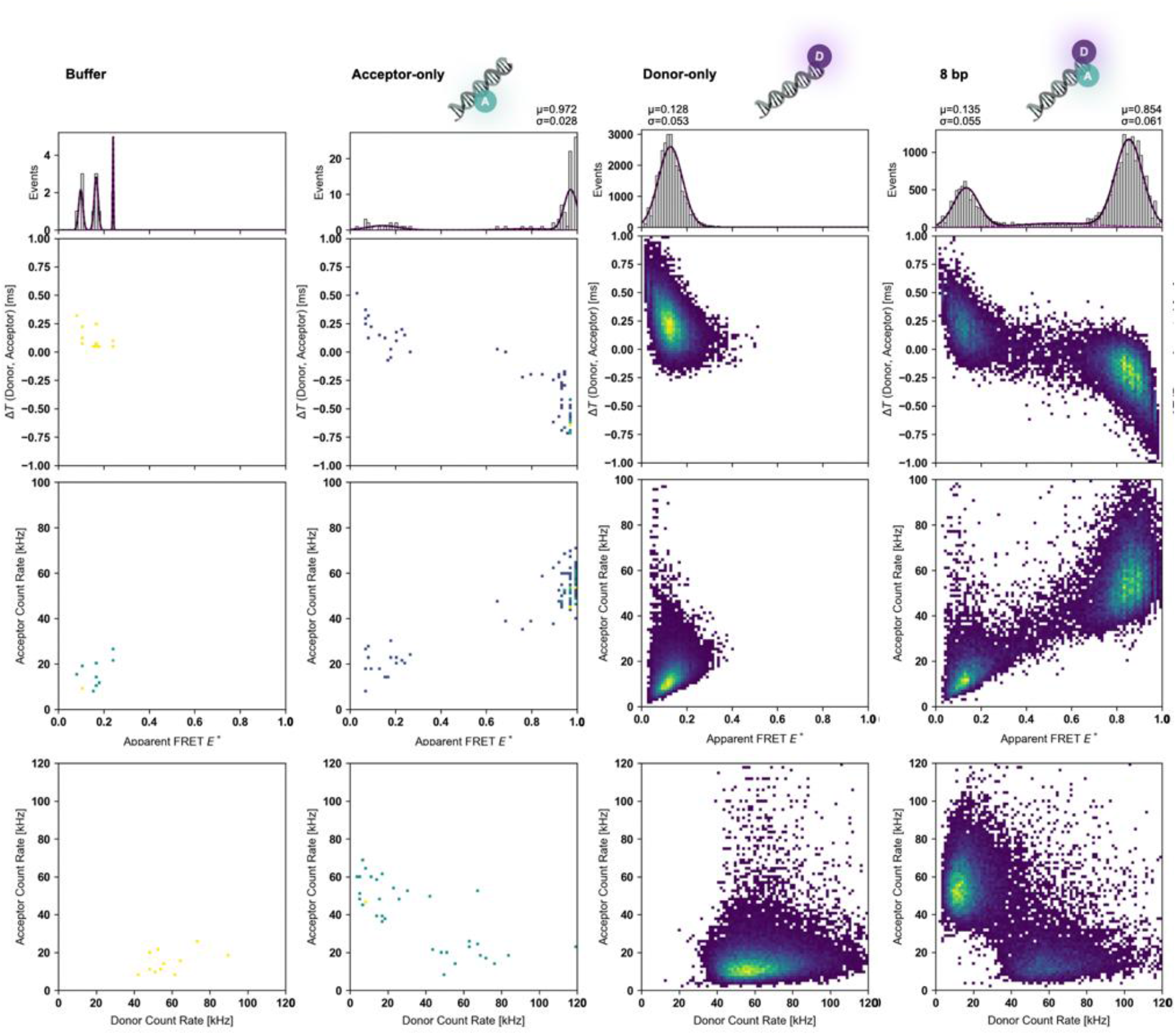
Supporting multiparameter 2D histograms of Alexa 488–Cy3B labeled dsDNA samples. Two-dimensional histograms of a 45-mer dsDNA labeled with Alexa 488 (donor) and Cy3B (acceptor) at 60 kW/cm^2^ excitation in the presence of 100 µM DAMF: PBS buffer, acceptor-only, donor-only and donor-acceptor with 8 bp interdye distances. Rows correspond to different parameter spaces: (top) apparent FRET efficiency versus ΔT_DA_, (middle) apparent FRET efficiency versus acceptor brightness, and (bottom) acceptor versus donor brightness.

**Figure S8:**
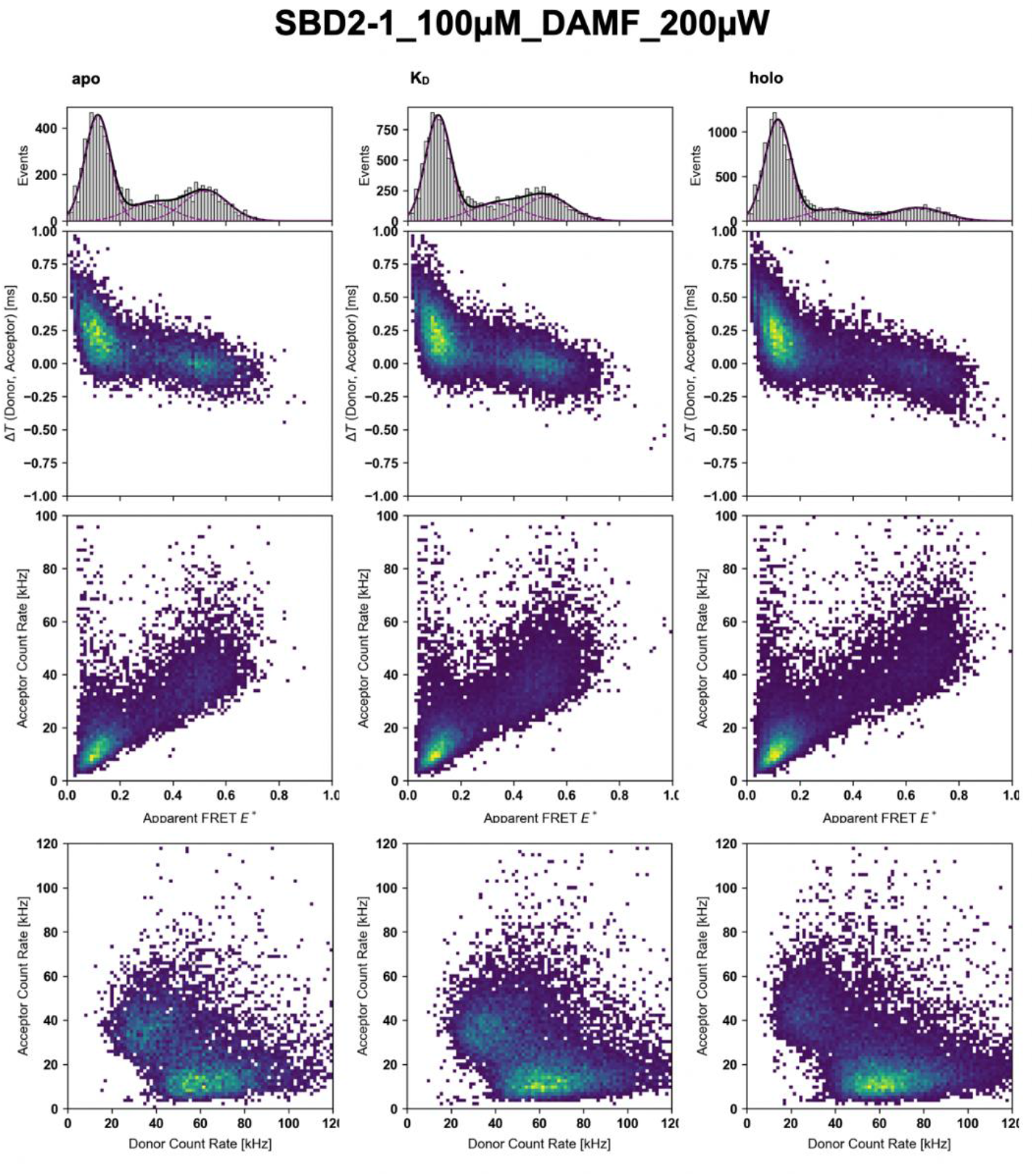
Multiparameter 2D histograms of SBD2 in different ligand-bound states with photostabilizer DAMF. ΔT-E, A-E and A-D histograms SBD2 with Alexa 488 (donor) and Cy3B (acceptor) at 60 kW/cm^2^ excitation with 100 µM DAMF. The left panel shows the apo (open), the middle and right panel shows the K_d_ (2 µM glutamine) conditions and holo state (1 mM glutamine), respectively.

**Figure S9:**
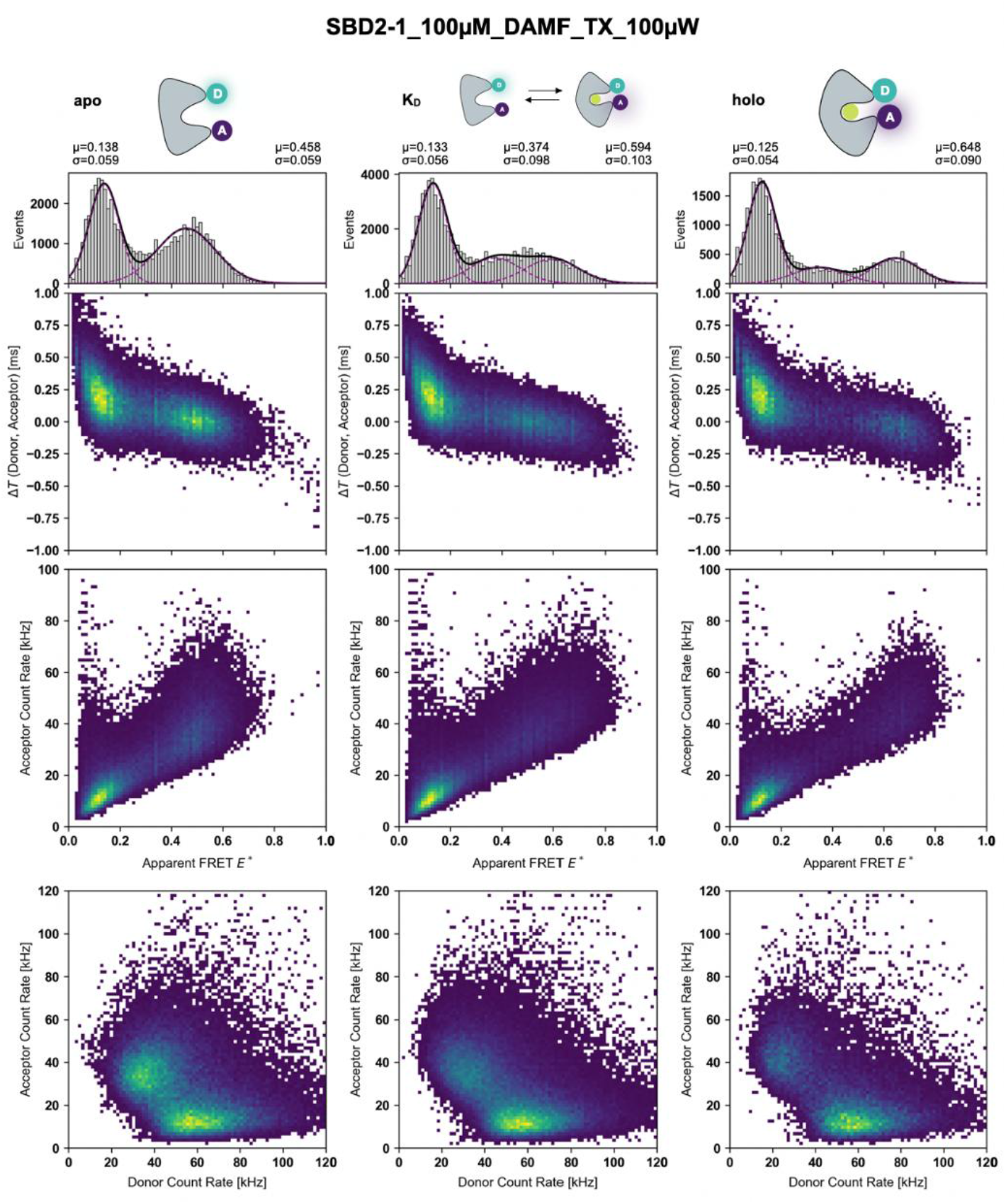
Multiparameter 2D histograms of SBD2 in different ligand-bound states with photostabilizer DAMF and Trolox. ΔT-E, A-E and A-D histograms SBD2 with Alexa 488 (donor) and Cy3B (acceptor) at 60 kW/cm^2^ excitation with 100 µM DAMF and 100 µM Trolox. The left panel shows the apo (open), the middle and right panel shows the K_d_ (2 µM glutamine) conditions and holo state (1 mM glutamine), respectively.

### Supplementary Note 1: Material and Methods

#### General sample preparation

##### Preparation of labeled DNA

Complementary single-strand fluorophore-labeled oligonucleotides^[1]^ were obtained from Ella Biotech (Fürstenfeldbruck, Germany) with either donor at the 5′-end of the top-strand (Alexa Fluor 488) or acceptor at different positions on the bottom strand (Alexa Fluor 555: 8/18, Cy3b: 8/18 or ATTO 542: 8/18/23 bp; table 1). Double-strand DNAs (dsDNA) were obtained by annealing using the following protocol: A 100-μl solution of two complementary single strand DNA (ssDNA) at a concentration of 1 μM was heated to 95°C for 4 min and then cooled down to 4°C at a rate of 1°C/min in an annealing buffer (20 mM Tris-HCl pH 8.0, 500 mM sodium chloride and 1 mM EDTA). The samples were kept at -20 °C until further usage.

**S2 Table :**
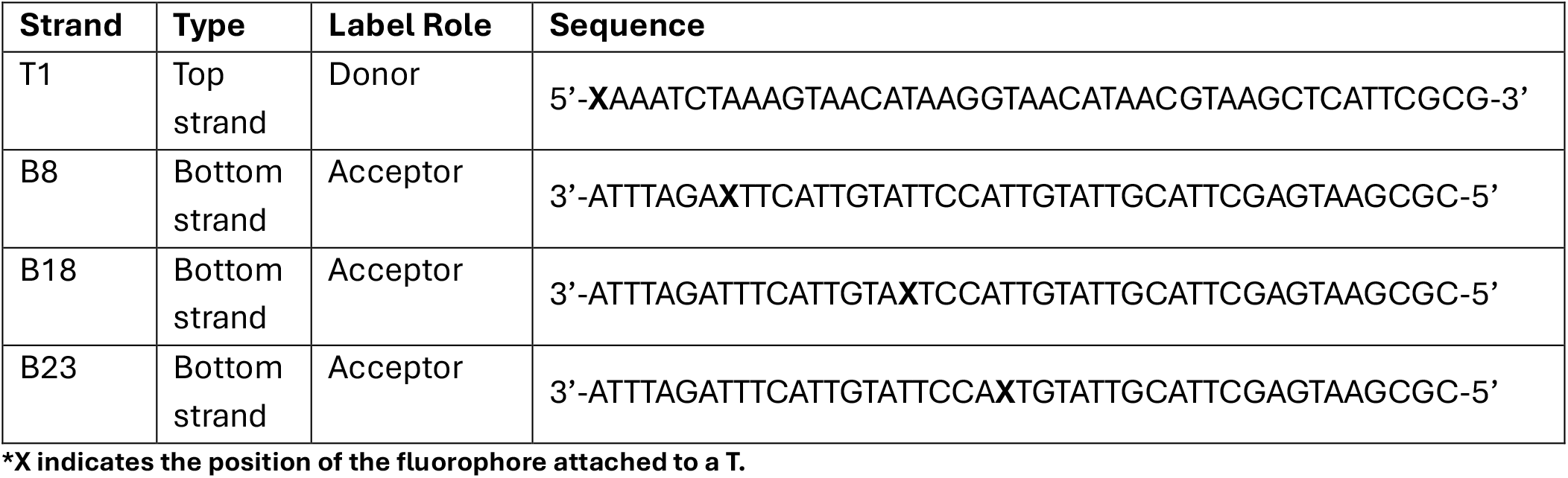
Sequences of DNA strands used in the study.

##### Protein expression and purification

SBD2 double-cysteine variant (T369C-S451C) was expressed and purified as previously described^[2]^. Briefly, a plasmid containing the coding region, including the appropriate mutations, was transformed into *Escherichia coli* BL21 (DE3) pLysS cells. The transformed cells were grown in 1 liter of LB medium containing 100 μg/mL Kanamycin and 50 μg/mL chloramphenicol at 37 °C under aerobic conditions. Bacterial growth was tracked using absorption measurements at 600 nm wavelength (OD_600nm_). Overexpression of the protein was induced at an OD_600nm_ of 0.6-0.7 by adding 1 mM IPTG to the culture media, followed by further incubation for 1.5-2.0 hours. Cells were centrifuged at 5000 x g for 30 min at 4 °C (Beckman, JA10). The cell pellet was resuspended in lysis buffer (50 mM Tris-HCl, pH 8.0, 1 M KCl, 10 mM imidazole, 10% glycerol and 1 mM dithiothreitol (DTT)) containing EDTA-free Protease Inhibitor Cocktail (cOmplete™, Roche), and incubated at 4 °C for 1 hour at gentle shaking. Following incubation the cells were disrupted by sonication (Branson tip sonication; amplitude: 25%; 10 min; 0.5 s on-off pulses; tube was kept in ice-water bath to avoid heating) and cell lysate was fractionated by two-step centrifugation (at 4 °C for 30 min at 4,416 g, Eppendorf, Centrifuge 5804 R) and at 4 °C for 1 hour for ultracentrifugation (70,658 g, Beckman, Type 70Ti under vacuum). The protein was purified by affinity chromatography using the Ni^2+^-Sepharose fast flow resin (GE Healthcare). The supernatant was loaded onto a pre-equilibrated resin and incubated for 1 hour at 4 °C with gentle shaking (resin equilibration included a 10 mL water wash followed by a 40 mL wash with lysis buffer).

The resin-bound protein was washed with 40 ml lysis buffer, followed by 40 ml of wash buffer I (50 mM Tris-HCl, pH 8,0, 50 mM KCl, 20 mM imidazole, 10% glycerol and 1 mM DTT), and finally eluted with 10 ml of elution buffer (50 mM Tris-HCl, pH 8.0, 50 mM KCl, 250 mM imidazole, 10% glycerol and 1 mM DTT). The eluted protein was dialyzed overnight against imidazole-free elution buffer, using Snakeskin^™^ dialysis membrane, to prevent unfolding/refolding interruptions. The dialyzed sample absorbance was measured at 280 nm, and the sample was aliquoted and stored at -80 °C until further handling.

##### Unfolding and refolding process of SBD2 (T369C-S451C)

The protein was diluted to a concentration of 4 µM in 50 ml of unfolding buffer (10 mM Hepes, pH 7.3 and 6M guanidine hydrochloride) and incubated for 3 hours at 30 °C with gentle shaking. Then, the sample was cooled down and centrifuged (3,046 x g for 30 minutes at 4 ℃) to remove insoluble aggregates prior to the refolding step. The supernatant was transferred to a Snakeskin^™^ dialysis membrane and dialyzed first against 2 l of L-arginine buffer (200 mM L-arginine, 150 mM NaCl, 10 mM Hepes pH 7.3, 5 mM DTT) for 18 hours, followed with 24 hours dialysis against 5 l dialysis buffer (150 mM NaCl, 10 mM Hepes pH 7.3, 5 mM DTT) with gentle stirring to ensure slow refolding and homogeneity. The refolded protein was then concentrated from 50 mL to a final 500 μL (Vivaspin 10 kDa MWCO; 3,000 g at 4 °C) and further purified by size-exclusion chromatography (ÄKTA pure system, Superdex-75 Increase 10/300 GL, GE Healthcare) and stored at -80 °C with 5% glycerol until further handling.

##### Protein labeling

The refolded SBD2 (T369C-S451C) was labelled as previously published^[2,3]^ using Alexa Fluor 488, Alexa Fluor 555 and Cy3B. Briefly, 0.6 mg of SBD2 (T369C-S451C) were incubated with 10 mM DTT ion labelling buffer (50 mM Tris-HCl pH 7.6, 150 mM NaCl) for 1 hour at 4 ℃. After incubation, the DTT concentration was reduced to 5 mM, and the protein sample was immobilized on 200 µL of Ni^2+^-Sepharose resin that had been washed and pre-equilibrated with labeling buffer. The resin was washed with 36 ml of labelling buffer followed by an overnight incubation with the fluorophores solution at 4 ℃ (50nmol of each fluorophore, dissolved in 0.7 ml of labelling buffer). Following incubation, excess unreacted fluorophores were removed by washing with 12 ml of labeling buffer, and the protein was eluted with 0.5 ml of labeling buffer containing 500 mM imidazole. The labelled protein was then purified by anion-exchange chromatography (AIEX) on an ÄKTA pure chromatography system (Cytiva; MonoQ 5/50 GL column, Cytiva) as previously described^[4]^.

### FCS-Guided Photostabilizer Screening Photostabilizers

#### Sample handling

DNA duplexes containing a single fluorophore were diluted to 0.5–1 nM in 100 µL PBS and measured on coverslips passivated with 1 mg/mL BSA in a series of 5–10 measurements each lasting 1 min. Photostabilizers were added to final concentrations of 1 mM Trolox ((±)-6-Hydroxy-2,5,7,8-tetramethylchromane-2-carboxylic acid, Sigma-Aldrich) or 100 µM DAMF ((Dimethylaminomethyl)ferrocen, Sigma-Aldrich).

#### 485-nm excitation (FCS of Alexa Fluor 488)

FCS experiments of Alexa Fluor 488 labeled dsDNA were conducted on an inverted microscope (MicroTime 200, PicoQuant) equipped with single photon counting electronics and picosecond time resolution (Hydra Harp 400, PicoQuant) or the FRET-Brick (see below). The sample was excited either through a 60x water immersion objective (Nikon M Plan Apo NA 1.20) or a 60x oil immersion objective (Olympus UPlanSApo NA 1.2) to in a diffraction-limited spo. The light emitted by the sample was collected through the same objective and split into perpendicular/parallel (polarizing beam splitter, PBS) components and detected by single-photon avalanche detectors (SPCM-AQR-14, Perkin Elmer) through green (BrightLine HC 520/35) band-pass filters. A linearly polarized laser diode (LDH-D-C-485 with 485 nm, PicoQuant) operated in continuous-wave mode excited the sample.

#### 532-nm excitation (FCS of Atto 542 and Cy3B)

For FCS experiments in the green spectral range, we used a custom-made ALEX microscope^[3]^. An OBIS 532-100-LS laser (Coherent, USA) provided continuous-wave excitation. The laser beam is coupled into a polarization-maintaining single-mode fiber (P3-57-488PM-FC-2, Thorlabs) via an aspheric fiber port (PAF2S-11A, Thorlabs), collimated (RC12APC-P01, Thorlabs) and guided into the epi-illuminated confocal microscope (Olympus IX71, Hamburg, Germany) by dual-edge beamsplitter ZT532/640rpc (Chroma/AHF) focused by a water immersion objective (UPlanSApo 60x/1.2w, Olympus Hamburg, Germany). The emitted fluorescence is collected through the objective and spatially filtered using a pinhole with 50 µm diameter and spectrally split into donor and acceptor channel by a single-edge dichroic mirror H643 LPXR (AHF). Fluorescence emission was filtered (donor: BrightLine HC 582/75 (Semrock/AHF), acceptor: Longpass 647 LP Edge Basic (Semrock/AHF), focused on avalanche photodiodes (SPCMAQRH-64, Excelitas). The detector outputs were recorded by a counter/timer device module (USB-CTR04, Measurement Computing, USA) using custom-made acquisition software written in Python, available as a compiled executable or editable code at https://github.com/harripd/mcc-daq-acquisition.

### The FRET-Brick

#### Optical configuration for the FRET-Brick (see also Figure 1a, main text)

For the excitation pathway, continuous-wave (CW) excitation was supplied by a USB-powered blue laser diode (488-30-1235-BL, Q-LINE). The laser power was adjusted through a continuous neutral density filter wheel (NDC-50C-2M, Thorlabs) and guided into an inversely mounted reflective collimator (RC08FC-P01, Thorlabs, USA), which coupled the beam into a polarization-maintaining single-mode fiber P3-57 488PM-FC-2 (Thorlabs, USA). The fiber guided the beam into the excitation layer, where it was collimated (RC12APC-P01, Thorlabs, USA). A dichroic beamsplitter with high reflectivity at 488 nm (ZT491rdc, Chroma/AHF, Germany) separated the excitation and emission beams to and from a high-NA apochromatic objective (60×, NA 1.2, UPlanSAPO 60XW, Olympus, Japan). For the emission and detection pathway, the emitted fluorescence was collected by the same objective and directed via a mirror into a piezo-directed optical mount (AG-M100N, Newport). The beam passed through an inversely mounted 12-mm reflective collimator (RC12FC-P01, Thorlabs), which focused and coupled the emission into a multimode optical fiber (10-µm core diameter, M64L01, Thorlabs). The fiber delivered the emission to a detection box, where it was collimated by a fixed-focus collimator (F220FC-532, Thorlabs) and spectrally split into two photon streams by a dichroic mirror (ZT543rdc longpass, Chroma/AHF, Germany). Individual photon streams were filtered with a band-pass filter for the donor channel (FF03-525/50, Semrock, Rochester NY, USA) and, for the acceptor channel, a 488-nm notch filter (NF488-15, Thorlabs, USA), and detected by PMTs (H10682-210, Hamamatsu, Japan). The detector outputs for FCS analysis were recorded via a counter/timer device module (USB-CTR04, Measurement Computing, USA) using custom-made acquisition software written in Python, available as a compiled executable or editable code at https://github.com/harripd/mcc-daq-acquisition.

#### 3D printing and assembly of the FRET-Brick

The FRET-Brick was assembled using components from previously printed parts for the µFCS and µALEX modality of the Brick-MIC^[4]^, which required no new parts to be printed. Specifically, the excitation layer of the µALEX modality was combined with the emission layer and detection box of the µFCS modality. All models were described in detail in ref. ^[4]^, which was designed and conceived using Onshape versions 1.114 to 1.172. 3D printing was carried out using PLATech filament (OLYMPfila) on an Ultimaker +2 Extended fitted with a 0.4-mm nozzle. All models were printed with an infill density of 17%, three layers for outer walls and with a layer height of 0.1 mm. The printing speed was set to 50 mm/s, and the nozzle temperature was maintained at 210°C. To prevent warping, all parts were printed with a brim and without any supports.

#### Sample handling

FRET-labeled 45-mer dsDNA samples were measured at a concentration of 100 pM in 100 µl droplets placed on pre-passivated coverslips (1 mg/ml BSA in PBS buffer), with or without the addition of the respective photostabilizers as detailed in the text and figure captions. The sample was excited at 488 nm with an excitation power of 200 µW (measured before entering the objective). Each measurement lasted on average 30 minutes.

Protein samples were measured at 100 pM in 100 µl droplets of 50 mM Tris-HCl pH 7.6, 150 mM NaCl buffer on pre-passivated coverslips (1 mg/ml BSA in PBS buffer). The sample was excited at 488 nm with an excitation power of 100 µW (measured before entering the objective). Proteins were recorded under three conditions: without ligand (apo), saturated with 1 mM glutamine (holo), and at an intermediate ligand concentration of 2 µM glutamine (approximately K_d_). Each condition was measured with 100 µM DAMF and 100 µM Trolox. Measurements lasted on average 45 minutes.

**Figure S10:**
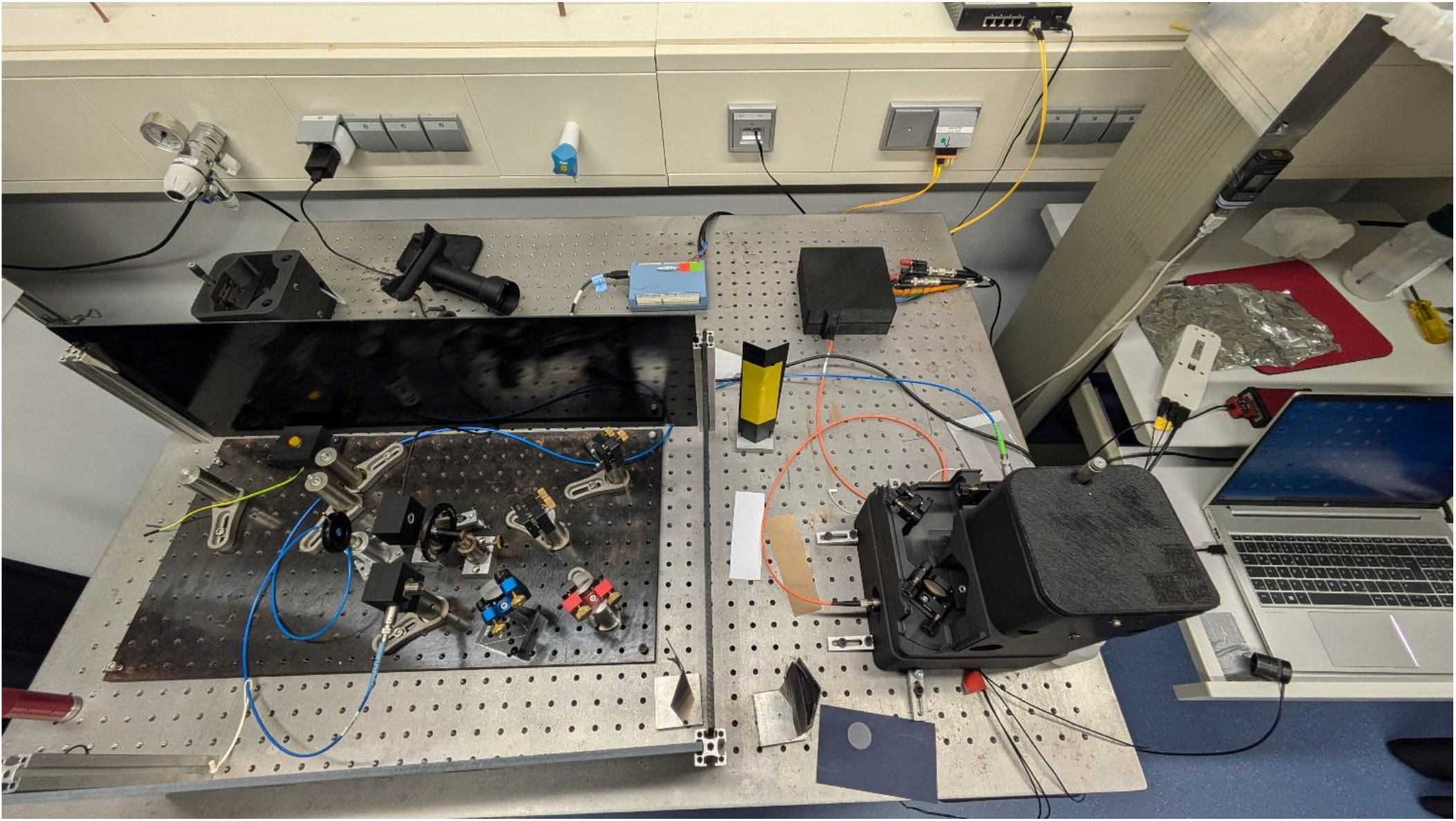
Photograph of the FRET-Brick.

### Supplementary Note 2: Single-molecule data analysis

#### Burst analysis

The used approaches to analyze photons within a burst (burst-wise analysis) groups are implemented in freely available open-source software^[5,6]^. Briefly, for every burst we determine spectroscopic parameters, such as channel, *X*, dependent integrated photon counts, *n*_*X*_, corresponding signal intensities, *S*_*X*_, apparent FRET efficiency E* and background, cross-talk and sensitivity-corrected FRET efficiencies, *E*,^[7]^ and the average burst pixel arrival times, *T*_*X*_ . We determine these parameters for different emission channels: referred to as *G* refers to the “blue” and *R* refers to “green”. The subscript “…|G” indicates excitation of the sample with a blue light source. The blue and green detection channels are denoted by the subscripts G|…, and R|…. The subscript “R|G” denotes green detection following blue excitation. The molecular species are indicated by superscripts; for instance, (DA) refers to a species. In the burst-wise single-molecule analysis, we group the photon stream into individual bursts by discriminating the background (approximately 1-2 kHz) from the fluorescence signal by applying an intensity threshold criterion^[8]^. Here, we use the total signal recorded for G|G and R|G to discriminate photons from the background. The background, the detection efficiency-ratio of the “blue” and “green” detectors, and the spectra were considered to determine the FRET efficiency, *E*, of every burst^[7]^. Following the grouping of photons, we determine for every photon group/pixel the signal intensities *S*_*B*|*B*_ and *S*_*G*|*B*_ to compute an apparent FRET efficiency E*:

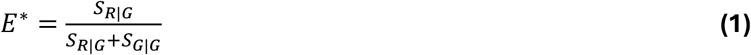

In addition to E* we compute *ΔT*(*G, R*), the difference between the duration of a burst when only inspecting *G*|*G* or *R*|*G* photons:

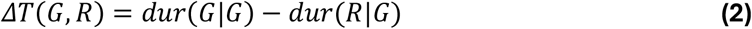

This additional burst metric makes use of time and channel information and identifies bursts, where the acceptor bleached within the burst duration.

#### Photon distribution analysis

To maximize the analysis precision of our single molecule experiments by explicitly considering the photon shot-noise of the data we used a Photon Distribution Analysis^[9]^ model implemented in the software ChiSurf^[10]^. The used model explicitly accounts for FRET, excitation cross talks (direct acceptor excitation), detector efficiencies, spectral crosstalks, and background contributions. Heterogeneities that are likely associated acceptor-induced histogram broadening beyond the shot-noise^[11]^ was modelled by a mixture of Gaussian distance distributions, which mapped to a distribution of FRET efficiencies. The parameters describing the fluorescence quantum yield and spectra were determined separately.

Briefly, the donor-acceptor distance distribution was modeled as normalized mixture of *K* Gaussians.

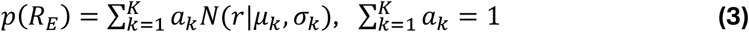

Where *R*_*E*_ is a distance, associated to the FRET efficiency, *μ*_*k*_, *σ*_*k*_, and *a*_*k*_ denote mean distance, width and amplitude of the k-th component, respectively. The corresponding FRE efficiency was obtained via:

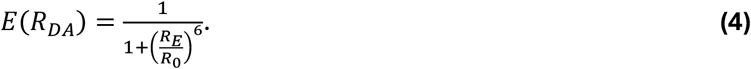

The distance distribution was discretized on a linear distance scale from 0.5 to 15.0 nm, resulting in a discretized *p*(*E*_*i*_) that is used in later steps to compute model FRET histograms. To account for excitation and emission crosstalks we first compute the fluorescence, intensity matrix, **I**

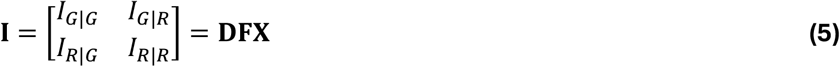

Where **X** is the excitation matrix dependent onf the light source, *L*

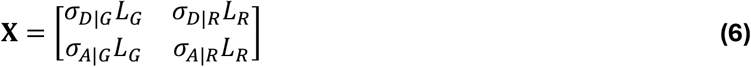

and *σ*_*d*|*l*_*L*_*s*_, the absorption of the fluorophore *d* ∈ {*D, A*} excited by the light source *s* ∈ {*R, G*}. Here, we performed OCE. Hence, *L*_*R*_=0. In **X**, off-diagonal elements are excitation cross-talks. Depending on the FRET efficiency, *E*, the intensities are redistributed across the emission channels.

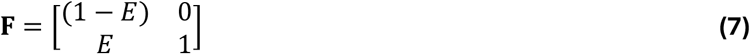

before being detected.

The detection is described by, **D**, the detection matrix that accounts for fluorescence quantum yields of the donor Φ_*F,D*_ and acceptor, Φ_*F,D*_, respectively,

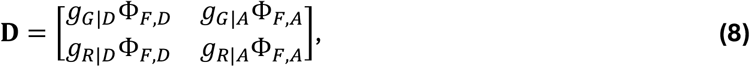

the spectral sensitivities *g*_*c*|*s*_ for the spectral channel, *c* ∈ {*G, R*}, and the species, *s* ∈ {*A, D*}. Off-diagonal elements are emission cross-talks.

Using the elements of the fluorescence intensity matrix, **I**, *p*_*G*_, the probability of the model for detecting a photon in the “Green” detection channel is:

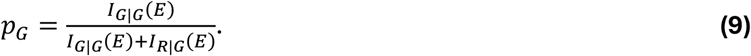

Given, *p*_*G*_, the conditional probability of observing a particular combination of green and red fluorescence photons, *F*_*G*_ and *F*_*R*_ probability, *P*(*F*_*G*_, *F*_*R*_ |*F*), for *F* registered fluorescence photons follows a binomial distribution:

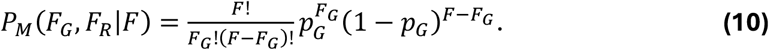

Here, we accounted for the background by modelling the background signals *B*_*G*_ and *B*_*R*_ by Poisson distributions, *P*(*B*_*G*_) and *P*(*B*_*R*_) to obtain the probability for a particular combination of fluorescent photons, *P*_*M*_(*S*_*G*_, *S*_*R*_):

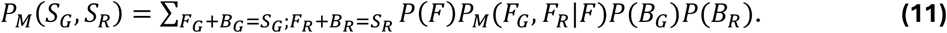

*P*(*F*) is the fluorescence intensity distribution, that is obtained from the total signal intensity distribution *P*(*S*) . To optimize model parameters such as *a*_*k*_, *μ*_*k*_ or *σ*_*k*_ we either score the modelled *P*_*M*_(*S*_*G*_, *S*_*R*_) or *P*_*M*_(*S*_*G*_, *S*_*R*_) marginalized to *E*_*app*_ against experimental histograms. The analysis procedure was implemented in the open-source software ChiSurf (https://github.com/fluorescence-tools/chisurf).

#### Fluorescence correlation spectroscopy (FCS)

Data analysis: Photon streams were binned to generate intensity traces and autocorrelated with a multi-tau algorithm^[12]^ implemented in the software Chisurf^[10]^ (https://github.com/fluorescence-tools/chisurf). The obtained autocorrelation functions, *G*(*τ*), were fitted to a 3D Brownian diffusion model:

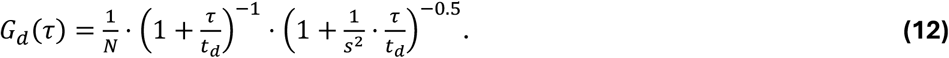

Here, *N, τ, t*_*d*_, and *s* are the effective number of molecules, the correlation time, the diffusion time, and a structure factor of the confocal volume, respectively.

Alexa Fluor 488 was described by a single dark-state relaxation term:

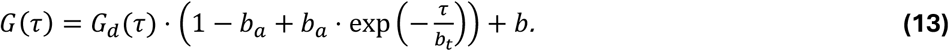

Experiments with Atto 542 and Cy3B were described by two dark-state relaxation terms:

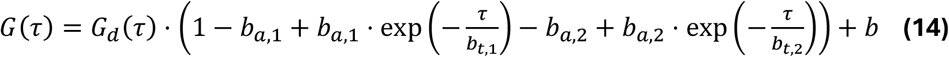

Above, *b, b*_*a*_, *b*_*t*_, are baseline offset, triplet amplitude, triplet lifetime, diffusion time, structure parameter, number of molecules and triplet relaxation term respectively.

## Footnotes

1 We note here that the CPM values reported in the FCS experiments in Figure 2 and Figure S6 were recorded on a Microtime 200 equipped with APD detectors. This implies that the reported CPM values are not the same as from the FRET-Brick which uses PMT detection (shown in Figure 1e and Figure S4). To allow a comparison of both setups we conducted comparative FCS experiment on both setups which revealed an approximate 5-10-fold brightness difference attributed to the sensitivity difference of APDs and PMTs (Table S1 vs. Figure S6).

## REFERENCES

[1] G. Agam, C. Gebhardt, M. Popara, R. Mächtel, J. Folz, B. Ambrose, N. Chamachi, S. Y. Chung, T. D. Craggs, M. de Boer, D. Grohmann, T. Ha, A. Hartmann, J. Hendrix, V. Hirschfeld, C. G. Hübner, T. Hugel, D. Kammerer, H.-S. Kang, A. N. Kapanidis, G. Krainer, K. Kramm, E. A. Lemke, E. Lerner, E. Margeat, K. Martens, J. Michaelis, J. Mitra, G. G. Moya Muñoz, R. B. Quast, N. C. Robb, M. Sattler, M. Schlierf, J. Schneider, T. Schröder, A. Sefer, P. S. Tan, J. Thurn, P. Tinnefeld, J. van Noort, S. Weiss, N. Wendler, N. Zijlstra, A. Barth, C. A. M. Seidel, D. C. Lamb, T. Cordes, “Reliability and accuracy of single-molecule FRET studies for characterization of structural dynamics and distances in proteins” Nat Methods 2023, 20, 523–535.

[2] B. Hellenkamp, S. Schmid, O. Doroshenko, O. Opanasyuk, R. Kühnemuth, S. Rezaei Adariani, B. Ambrose, M. Aznauryan, A. Barth, V. Birkedal, M. E. Bowen, H. Chen, T. Cordes, T. Eilert, C. Fijen, C. Gebhardt, M. Götz, G. Gouridis, E. Gratton, T. Ha, P. Hao, C. A. Hanke, A. Hartmann, J. Hendrix, L. L. Hildebrandt, V. Hirschfeld, J. Hohlbein, B. Hua, C. G. Hübner, E. Kallis, A. N. Kapanidis, J.-Y. Kim, G. Krainer, D. C. Lamb, N. K. Lee, E. A. Lemke, B. Levesque, M. Levitus, J. J. McCann, N. Naredi-Rainer, D. Nettels, T. Ngo, R. Qiu, N. C. Robb, C. Röcker, H. Sanabria, M. Schlierf, T. Schröder, B. Schuler, H. Seidel, L. Streit, J. Thurn, P. Tinnefeld, S. Tyagi, N. Vandenberk, A. M. Vera, K. R. Weninger, B. Wünsch, I. S. Yanez-Orozco, J. Michaelis, C. A. M. Seidel, T. D. Craggs, T. Hugel, “Precision and accuracy of single-molecule FRET measurements—a multilaboratory benchmark study” Nat Methods 2018, 15, 669–676.

[3] E. Sisamakis, A. Valeri, S. Kalinin, P. J. Rothwell, C. A. M. Seidel, “Accurate single-molecule FRET studies using multiparameter fluorescence detection” Methods Enzymol 2010, 475, 455–514.

[4] E. Lerner, T. Cordes, A. Ingargiola, Y. Alhadid, S. Chung, X. Michalet, S. Weiss, “Toward dynamic structural biology: Two decades of single-molecule Förster resonance energy transfer” Science 2018, 359, eaan1133.

[5] E. Lerner, A. Barth, J. Hendrix, B. Ambrose, V. Birkedal, S. C. Blanchard, R. Börner, H. Sung Chung, T. Cordes, T. D. Craggs, A. A. Deniz, J. Diao, J. Fei, R. L. Gonzalez, I. V. Gopich, T. Ha, C. A. Hanke, G. Haran, N. S. Hatzakis, S. Hohng, S.-C. Hong, T. Hugel, A. Ingargiola, C. Joo, A. N. Kapanidis, H. D. Kim, T. Laurence, N. K. Lee, T.-H. Lee, E. A. Lemke, E. Margeat, J. Michaelis, X. Michalet, S. Myong, D. Nettels, T.-O. Peulen, E. Ploetz, Y. Razvag, N. C. Robb, B. Schuler, H. Soleimaninejad, C. Tang, R. Vafabakhsh, D. C. Lamb, C. A. Seidel, S. Weiss, “FRET-based dynamic structural biology: Challenges, perspectives and an appeal for open-science practices” eLife n.d., 10, e60416.

[6] N. K. Lee, A. N. Kapanidis, Y. Wang, X. Michalet, J. Mukhopadhyay, R. H. Ebright, S. Weiss, “Accurate FRET Measurements within Single Diffusing Biomolecules Using Alternating-Laser Excitation” Biophysical Journal 2005, 88, 2939–2953.

[7] A. Muschielok, J. Andrecka, A. Jawhari, F. Brückner, P. Cramer, J. Michaelis, “A nano-positioning system for macromolecular structural analysis” Nat Methods 2008, 5, 965–971.

[8] S. Kalinin, T. Peulen, S. Sindbert, P. J. Rothwell, S. Berger, T. Restle, R. S. Goody, H. Gohlke, C. A. M. Seidel, “A toolkit and benchmark study for FRET-restrained high-precision structural modeling” Nat Methods 2012, 9, 1218–1225.

[9] B. Hellenkamp, P. Wortmann, F. Kandzia, M. Zacharias, T. Hugel, “Multidomain structure and correlated dynamics determined by self-consistent FRET networks” Nat Methods 2017, 14, 174–180.

[10] T. D. Craggs, A. N. Kapanidis, “Six steps closer to FRET-driven structural biology” Nat Methods 2012, 9, 1157–1158.

[11] M. Dimura, T. O. Peulen, C. A. Hanke, A. Prakash, H. Gohlke, C. A. Seidel, “Quantitative FRET studies and integrative modeling unravel the structure and dynamics of biomolecular systems” Curr Opin Struct Biol 2016, 40, 163–185.

[12] J. Hohlbein, T. D. Craggs, T. Cordes, “Alternating-laser excitation: single-molecule FRET and beyond” Chem. Soc. Rev. 2014, 43, 1156–1171.

[13] B. K. Müller, E. Zaychikov, C. Bräuchle, D. C. Lamb, “Pulsed Interleaved Excitation” Biophysical Journal 2005, 89, 3508–3522.

[14] E. Bilgen, D. C. Lamb, “Multicolor single-molecule FRET studies on dynamic protein systems” Current Opinion in Structural Biology 2025, 93, 103117.

[15] L. Bonhomme, E. Bilgen, C. Clerté, J.-P. Pin, P. Rondard, E. Margeat, D. C. Lamb, R. B. Quast, “Triple Labeling Resolves a GPCR Intermediate State by Using Three-Color Single Molecule FRET” J. Am. Chem. Soc. 2025, 147, 17689–17700.

[16] J. Widengren, V. Kudryavtsev, M. Antonik, S. Berger, M. Gerken, C. A. M. Seidel, “Single-Molecule Detection and Identification of Multiple Species by Multiparameter Fluorescence Detection” Anal. Chem. 2006, 78, 2039–2050.

[17] P. J. Rothwell, S. Berger, O. Kensch, S. Felekyan, M. Antonik, B. M. Wöhrl, T. Restle, R. S. Goody, C. A. M. Seidel, “Multiparameter single-molecule fluorescence spectroscopy reveals heterogeneity of HIV-1 reverse transcriptase:primer/template complexes” Proc. Natl. Acad. Sci. U.S.A. 2003, 100, 1655–1660.

[18] A. N. Kapanidis, N. K. Lee, T. A. Laurence, S. Doose, E. Margeat, S. Weiss, “Fluorescence-aided molecule sorting: analysis of structure and interactions by alternating-laser excitation of single molecules” Proc Natl Acad Sci U S A 2004, 101, 8936–8941.

[19] A. N. Kapanidis, T. A. Laurence, N. K. Lee, E. Margeat, X. Kong, S. Weiss, “Alternating-Laser Excitation of Single Molecules” Acc. Chem. Res. 2005, 38, 523–533.

[20] F. Huang, S. Sato, T. D. Sharpe, L. Ying, A. R. Fersht, “Distinguishing between cooperative and unimodal downhill protein folding” Proc. Natl. Acad. Sci. U.S.A. 2007, 104, 123–127.

[21] F. Husada, K. Bountra, K. Tassis, M. De Boer, M. Romano, S. Rebuffat, K. Beis, T. Cordes, “Conformational dynamics of the ABC transporter McjD seen by single-molecule FRET” The EMBO Journal 2018, 37, e100056.

[22] D. Scheerer, D. Levy, R. Casier, I. Riven, H. Mazal, G. Haran, “Interplay between conformational dynamics and substrate binding regulates enzymatic activity: a single-molecule FRET study” Chem. Sci. 2025, 16, 3066–3077.

[23] D. S. Majumdar, I. Smirnova, V. Kasho, E. Nir, X. Kong, S. Weiss, H. R. Kaback, “Single-molecule FRET reveals sugar-induced conformational dynamics in LacY” Proc. Natl. Acad. Sci. U.S.A. 2007, 104, 12640–12645.

[24] J. Fort, A. Nicolàs-Aragó, L. Maggi, M. Martinez-Molledo, D. Kapiki, P. González-Novoa, P. Gómez-Gejo, N. Zijlstra, S. Bodoy, E. Pardon, J. Steyaert, O. Llorca, M. Orozco, T. Cordes, M. Palacín, “The conserved lysine residue in transmembrane helix 5 is pivotal for the cytoplasmic gating of the Lamino acid transporters” PNAS Nexus 2024, 4, pgae584.

[25] Y. Santoso, J. P. Torella, A. N. Kapanidis, “Characterizing Single-Molecule FRET Dynamics with Probability Distribution Analysis” ChemPhysChem 2010, 11, 2209–2219.

[26] M. de Boer, G. Gouridis, R. Vietrov, S. L. Begg, G. K. Schuurman-Wolters, F. Husada, N. Eleftheriadis, B. Poolman, C. A. McDevitt, T. Cordes, “Conformational and dynamic plasticity in substrate-binding proteins underlies selective transport in ABC importers” eLife 2019, 8, e44652.

[27] G. Gouridis, Y. A. Muthahari, M. De Boer, D. A. Griffith, A. Tsirigotaki, K. Tassis, N. Zijlstra, R. Xu, N. Eleftheriadis, Y. Sugijo, M. Zacharias, A. Dömling, S. Karamanou, C. Pozidis, A. Economou, T. Cordes, “Structural dynamics in the evolution of a bilobed protein scaffold” Proc. Natl. Acad. Sci. U.S.A. 2021, 118, e2026165118.

[28] E. Ploetz, G. K. Schuurman-Wolters, N. Zijlstra, A. W. Jager, D. A. Griffith, A. Guskov, G. Gouridis, B. Poolman, T. Cordes, “Structural and biophysical characterization of the tandem substrate-binding domains of the ABC importer GlnPQ” Open Biol. 2021, 11, 200406.

[29] G. G. Moya Muñoz, O. Brix, P. Klocke, P. D. Harris, J. R. Luna Piedra, N. D. Wendler, E. Lerner, N. Zijlstra, T. Cordes, “Single-molecule detection and super-resolution imaging with a portable and adaptable 3D-printed microscopy platform (Brick-MIC)” Sci. Adv. 2024, 10, eado3427.

[30] J. W. P. Brown, A. Bauer, M. E. Polinkovsky, A. Bhumkar, D. J. B. Hunter, K. Gaus, E. Sierecki, Y. Gambin, “Single-molecule detection on a portable 3D-printed microscope” Nat Commun 2019, 10, 5662.

[31] R. Strack, “The miCube open microscope” Nat Methods 2019, 16, 958.

[32] B. Ambrose, J. M. Baxter, J. Cully, M. Willmott, E. M. Steele, B. C. Bateman, M. L. Martin-Fernandez, A. Cadby, J. Shewring, M. Aaldering, T. D. Craggs, “The smfBox is an open-source platform for single-molecule FRET” Nat Commun 2020, 11, 5641.

[33] B. Diederich, R. Lachmann, S. Carlstedt, B. Marsikova, H. Wang, X. Uwurukundo, A. S. Mosig, R. Heintzmann, “A versatile and customizable low-cost 3D-printed open standard for microscopic imaging” Nat Commun 2020, 11, 5979.

[34] A. C. Zehrer, A. Martin-Villalba, B. Diederich, H. Ewers, 2024, DOI: 10.7554/eLife.89826.2.

[35] M. Loretan, M. Barella, N. Fuchs, S. Kocabey, K. Kołątaj, F. D. Stefani, G. P. Acuna, “Direct single-molecule detection and super-resolution imaging with a low-cost portable smartphone-based microscope” Nat Commun 2025, 16, 8937.

[36] J. Lightley, S. Kumar, M. Q. Lim, E. Garcia, F. Görlitz, Y. Alexandrov, T. Parrado, C. Hollick, E. Steele, K. Roßmann, J. Graham, J. Broichhagen, I. A. McNeish, C. A. Roufosse, M. A. A. Neil, C. Dunsby, P. M. W. French, “openFrame : A modular, sustainable, open microscopy platform with single-shot, dual-axis optical autofocus module providing high precision and long range of operation” Journal of Microscopy 2023, 292, 64–77.

[37] A. Ingargiola, T. Laurence, R. Boutelle, S. Weiss, X. Michalet, “Photon-HDF5: An Open File Format for Timestamp-Based Single-Molecule Fluorescence Experiments” Biophysical Journal 2016, 110, 26–33.

[38] T.-O. Peulen, O. Opanasyuk, C. A. M. Seidel, “Combining Graphical and Analytical Methods with Molecular Simulations To Analyze Time-Resolved FRET Measurements of Labeled Macromolecules Accurately” J. Phys. Chem. B 2017, 121, 8211–8241.

[39] H. Wallrabe, M. Stanley, A. Periasamy, M. Barroso, “One- and two-photon fluorescence resonance energy transfer microscopy to establish a clustered distribution of receptor-ligand complexes in endocytic membranes” J. Biomed. Opt. 2003, 8, 339.

[40] A. Konrad, M. Metzger, A. M. Kern, M. Brecht, A. J. Meixner, “Revealing the radiative and non-radiative relaxation rates of the fluorescent dye Atto488 in a λ/2 Fabry–Pérot-resonator by spectral and time resolved measurements” Nanoscale 2016, 8, 14541–14547.

[41] N. Panchuk-Voloshina, R. P. Haugland, J. Bishop-Stewart, M. K. Bhalgat, P. J. Millard, F. Mao, W.-Y. Leung, R. P. Haugland, “Alexa Dyes, a Series of New Fluorescent Dyes that Yield Exceptionally Bright, Photostable Conjugates” J Histochem Cytochem. 1999, 47, 1179–1188.

[42] I. Rasnik, S. A. McKinney, T. Ha, “Nonblinking and long-lasting single-molecule fluorescence imaging” Nat Methods 2006, 3, 891–893.

[43] T. Cordes, J. Vogelsang, P. Tinnefeld, “On the Mechanism of Trolox as Antiblinking and Antibleaching Reagent” J. Am. Chem. Soc. 2009, 131, 5018–5019.

[44] T. Cordes, I. H. Stein, C. Forthmann, C. Steinhauer, M. Walz, W. Summerer, B. Person, J. Vogelsang, P. Tinnefeld in Advanced Microscopy Techniques, OSA, Munich, 2009, p. 7367_1D.

[45] L. Zhang, C. Wang, Y. Li, H. Wang, K. Sun, S. Lu, Y. Wang, S. Jing, T. Cordes, “Modular Design and Scaffold-Synthesis of Multi-Functional Fluorophores for Targeted Cellular Imaging and Pyroptosis” Angew Chem Int Ed 2025, 64, e202415627.

[46] H. Niu, J. Liu, H. M. O’Connor, T. Gunnlaugsson, T. D. James, H. Zhang, “Photoinduced electron transfer (PeT) based fluorescent probes for cellular imaging and disease therapy” Chem. Soc. Rev. 2023, 52, 2322– 2357.

[47] J. Vogelsang, R. Kasper, C. Steinhauer, B. Person, M. Heilemann, M. Sauer, P. Tinnefeld, “A Reducing and Oxidizing System Minimizes Photobleaching and Blinking of Fluorescent Dyes” Angew Chem Int Ed 2008, 47, 5465–5469.

[48] G. Gouridis, G. K. Schuurman-Wolters, E. Ploetz, F. Husada, R. Vietrov, M. de Boer, T. Cordes, B. Poolman, “Conformational dynamics in substrate-binding domains influences transport in the ABC importer GlnPQ” Nat Struct Mol Biol 2015, 22, 57–64.

[49] L. Le Reste, J. Hohlbein, K. Gryte, A. N. Kapanidis, “Characterization of Dark Quencher Chromophores as Nonfluorescent Acceptors for Single-Molecule FRET” Biophysical Journal 2012, 102, 2658–2668.

[50] E. Ploetz, E. Lerner, F. Husada, M. Roelfs, S. Chung, J. Hohlbein, S. Weiss, T. Cordes, “Förster resonance energy transfer and protein-induced fluorescence enhancement as synergetic multi-scale molecular rulers” Sci Rep 2016, 6, 33257.

[51] R. P.-A. Berntsson, S. H. J. Smits, L. Schmitt, D.-J. Slotboom, B. Poolman, “A structural classification of substrate-binding proteins” FEBS Letters 2010, 584, 2606–2617.

[52] G. H. Scheepers, J. A. Lycklama A Nijeholt, B. Poolman, “An updated structural classification of substrate-binding proteins” FEBS Letters 2016, 590, 4393–4401.

[53] M. F. Peter, C. Gebhardt, R. Mächtel, G. G. M. Muñoz, J. Glaenzer, A. Narducci, G. H. Thomas, T. Cordes, G. Hagelueken, “Cross-validation of distance measurements in proteins by PELDOR/DEER and single-molecule FRET” Nat Commun 2022, 13, 4396.

[54] S. Kalinin, S. Felekyan, M. Antonik, C. A. M. Seidel, “Probability Distribution Analysis of Single-Molecule Fluorescence Anisotropy and Resonance Energy Transfer” J. Phys. Chem. B 2007, 111, 10253–10262.

[55] M. E. Sanborn, B. K. Connolly, K. Gurunathan, M. Levitus, “Fluorescence Properties and Photophysics of the Sulfoindocyanine Cy3 Linked Covalently to DNA” J. Phys. Chem. B 2007, 111, 11064–11074.

[56] T. J. Lambert, “FPbase: a community-editable fluorescent protein database” Nat Methods 2019, 16, 277– 278.

[57] X. A. Feng, M. F. Poyton, T. Ha, “Multicolor single-molecule FRET for DNA and RNA processes” Current Opinion in Structural Biology 2021, 70, 26–33.

[58] I. H. Stein, C. Steinhauer, P. Tinnefeld, “Single-Molecule Four-Color FRET Visualizes Energy-Transfer Paths on DNA Origami” J. Am. Chem. Soc. 2011, 133, 4193–4195.

[59] V. Kudryavtsev, M. Sikor, S. Kalinin, D. Mokranjac, C. A. M. Seidel, D. C. Lamb, “Combining MFD and PIE for Accurate Single-Pair Förster Resonance Energy Transfer Measurements” ChemPhysChem 2012, 13, 1060– 1078.

[60] T. A. Laurence, X. Kong, M. Jäger, S. Weiss, “Probing structural heterogeneities and fluctuations of nucleic acids and denatured proteins” Proc. Natl. Acad. Sci. U.S.A. 2005, 102, 17348–17353.

[61] A. Ingargiola, R. A. Colyer, D. Kim, F. Panzeri, R. Lin, A. Gulinatti, I. Rech, M. Ghioni, S. Weiss, X. Michalet in (Eds.: J. Enderlein, Z.K. Gryczynski, R. Erdmann, F. Koberling, I. Gregor), San Francisco, California, USA, 2012, p. 82280B.

[62] R. Kiselev, R. A. Brady, A. Modak, F. A. Cruz-Navarrete, J. L. Alejo, D. S. Terry, R. B. Altman, W. B. Asher, J. A. Javitch, S. C. Blanchard, “Parallel stopped-flow interrogation of diverse biological systems at the single-molecule scale” Nat Methods 2025, DOI 10.1038/s41592-025-02944-4.

[63] A. Hartmann, K. Sreenivasa, M. Schenkel, N. Chamachi, P. Schake, G. Krainer, M. Schlierf, “An automated single-molecule FRET platform for high-content, multiwell plate screening of biomolecular conformations and dynamics” Nat Commun 2023, 14, 6511.

[64] S. Kim, A. M. Streets, R. R. Lin, S. R. Quake, S. Weiss, D. S. Majumdar, “High-throughput single-molecule optofluidic analysis” Nat Methods 2011, 8, 242–245.

## SI References

[1] E. Ploetz, E. Lerner, F. Husada, M. Roelfs, S. Chung, J. Hohlbein, S. Weiss, T. Cordes, “Förster resonance energy transfer and protein-induced fluorescence enhancement as synergetic multi-scale molecular rulers” Sci Rep 2016, 6, 33257.

[2] G. Gouridis, G. K. Schuurman-Wolters, E. Ploetz, F. Husada, R. Vietrov, M. de Boer, T. Cordes, B. Poolman, “Conformational dynamics in substrate-binding domains influences transport in the ABC importer GlnPQ” Nat Struct Mol Biol 2015, 22, 57–64.

[3] M. de Boer, G. Gouridis, R. Vietrov, S. L. Begg, G. K. Schuurman-Wolters, F. Husada, N. Eleftheriadis, B. Poolman, C. A. McDevitt, T. Cordes, “Conformational and dynamic plasticity in substrate-binding proteins underlies selective transport in ABC importers” eLife 2019, 8, e44652.

[4] G. G. Moya Muñoz, O. Brix, P. Klocke, P. D. Harris, J. R. Luna Piedra, N. D. Wendler, E. Lerner, N. Zijlstra, T. Cordes, “Single-molecule detection and super-resolution imaging with a portable and adaptable 3D-printed microscopy platform (Brick-MIC)” Sci. Adv. 2024, 10, eado3427.

[5] T.-O. Peulen, K. Hemmen, A. Greife, B. M. Webb, S. Felekyan, A. Sali, C. A. M. Seidel, H. Sanabria, K. G. Heinze, “tttrlib: modular software for integrating fluorescence spectroscopy, imaging, and molecular modeling” Bioinformatics 2025, 41, btaf025.

[6] T.-O. Peulen, O. Opanasyuk, C. A. M. Seidel, “Combining Graphical and Analytical Methods with Molecular Simulations To Analyze Time-Resolved FRET Measurements of Labeled Macromolecules Accurately” J. Phys. Chem. B 2017, 121, 8211–8241.

[7] E. Sisamakis, A. Valeri, S. Kalinin, P. J. Rothwell, C. A. M. Seidel, “Accurate single-molecule FRET studies using multiparameter fluorescence detection” Methods Enzymol 2010, 475, 455–514.

[8] C. Eggeling, S. Berger, L. Brand, J. R. Fries, J. Schaffer, A. Volkmer, C. A. M. Seidel, “Data registration and selective single-molecule analysis using multi-parameter fluorescence detection” Journal of Biotechnology 2001, 86, 163–180.

[9] S. Kalinin, S. Felekyan, M. Antonik, C. A. M. Seidel, “Probability Distribution Analysis of Single-Molecule Fluorescence Anisotropy and Resonance Energy Transfer” J. Phys. Chem. B 2007, 111, 10253–10262.

[10] T.-O. Peulen, “Exploring Time-Resolved Fluorescence Data: A Software Solution for Model Generation and Analysis” Spectroscopy Journal 2025, 3, 16.

[11] S. Kalinin, E. Sisamakis, S. W. Magennis, S. Felekyan, C. A. M. Seidel, “On the Origin of Broadening of Single-Molecule FRET Efficiency Distributions beyond Shot Noise Limits” J. Phys. Chem. B 2010, 114, 6197–6206.

[12] S. Felekyan, R. Kühnemuth, V. Kudryavtsev, C. Sandhagen, W. Becker, C. A. M. Seidel, “Full correlation from picoseconds to seconds by time-resolved and time-correlated single photon detection” Review of Scientific Instruments 2005, 76, 083104.

